# Specialized and spatially organized coding of sensory, motor, and cognitive variables in midbrain dopamine neurons

**DOI:** 10.1101/456194

**Authors:** Ben Engelhard, Joel Finkelstein, Julia Cox, Weston Fleming, Hee Jae Jang, Sharon Ornelas, Sue Ann Koay, Stephan Thiberge, Nathaniel Daw, David W. Tank, Ilana B. Witten

**Affiliations:** Princeton Neuroscience Institute, Princeton University, Princeton NJ 08544; Department of Psychology, Princeton University, Princeton NJ 08544; Department of Molecular Biology, Princeton University, Princeton NJ 08544

## Abstract

There is increased appreciation that dopamine (DA) neurons in the midbrain respond not only to reward ^1^^,^^2^ and reward-predicting cues ^1^^,^^3^^,^^4^^,^ but also to other variables such as distance to reward 5, movements ^6^^–^^11^ and behavioral choices 12–15. Based on these findings, a major open question is how the responses to these diverse variables are organized across the population of DA neurons. In other words, do individual DA neurons multiplex multiple variables, or are subsets of neurons specialized in encoding specific behavioral variables? The reason that this fundamental question has been difficult to resolve is that recordings from large populations of individual DA neurons have not been performed in a behavioral task with sufficient complexity to examine these diverse variables simultaneously. To address this gap, we used 2-photon calcium imaging through an implanted lens to record activity of >300 midbrain DA neurons in the VTA during a complex decision-making task. As mice navigated in a virtual reality (VR) environment, DA neurons encoded an array of sensory, motor, and cognitive variables. These responses were functionally clustered, such that subpopulations of neurons transmitted information about a subset of behavioral variables, in addition to encoding reward. These functional clusters were spatially organized, such that neighboring neurons were more likely to be part of the same cluster. Taken together with the topography between DA neurons and their projections, this specialization and anatomical organization may aid downstream circuits in correctly interpreting the wide range of signals transmitted by DA neurons.

---

To determine how responses are organized across the population of VTA DA neurons, we sought to record at cellular resolution from ensembles of identified DA neurons in a behavioral task with sufficient complexity to engage many of the behavioral variables that are now thought to be of relevance to DA neurons. These variables include reward ^1^^,^^2^^,^^16^^,^^17^, reward-predicting cues ^1^^,^^3^, reward history ^13^^,^^18^, spatial position ^5^, kinematics (velocity, acceleration, view angle) ^6^^–^^9^, and behavioral choices ^12^^,^^13^^,^^19^.

Towards this end, we trained 20 mice on a decision making task in a VR environment that encompassed this wide range of behavioral variables (“Accumulating Towers” task ^20^; Fig. 1a,b; visual snapshots of the maze in Extended Data Fig. 1a). As mice navigated the central stem of the virtual T-maze, they observed transient reward-predicting cues on the left and right of the maze stem that signaled which maze arm was most likely to be rewarded (“cue period”; Fig. 1b). By turning to the side of the maze with more cues, the mice received a water reward, while turning to the other side resulted in a tone and a 3s time out. The 2s period after reward delivery or tone presentation was termed the “outcome period” (Fig. 1b). As expected, after training, mice tended to turn to the maze arm associated with more cues (Fig. 1c; average percent correct is 77.6±0.9%).

**Figure 1.**
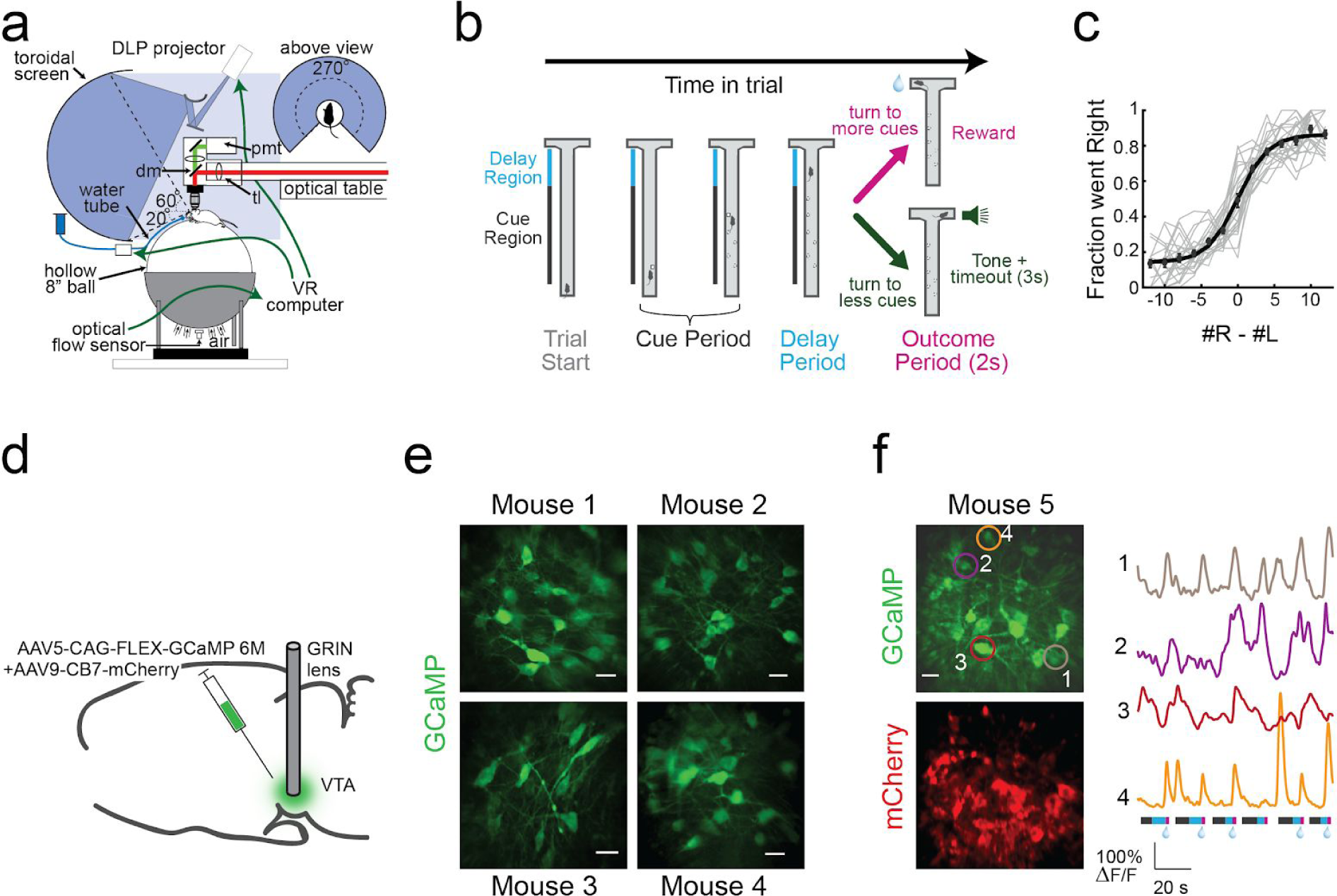
2-photon imaging of VTA DA neuron during navigation and decision making in virtual reality. **a,** Schematic of the behavioral and imaging setup. DLP: Digital Light Processing. pmt: photomultiplier <u>tube. dm:</u> dichroic mirror. tl: tube lens. **b,** Schematic of an example trial in the virtual T-maze. In the central stem of the maze, the mouse is presented with transient, Poisson-distributed visual cues to the left and right (“cue period”). At the end of the central stem, turning to the arm side with more cues results in reward delivery, while turning to the other arm results in a tone and a 3s timeout. **c,** Psychometric curves of all behavioral sessions that correspond to data in this paper. Gray lines are individual sessions. Circles and bars are mean±s.e.m. Black line is a logistic fit to the mean across sessions. **d,** Schematic of the surgical strategy. **e,** Fields of view through the GRIN lens for 4 example mice. The horizontal white bars are 20 um scale bars. **f,** Left: Simultaneously recorded fields of view through the green and red channels. Right: traces from 4 example neurons during 6 consecutive trials. The bars below the traces indicates the timing of each epoch within each trial: cue period (grey), delay period (blue), outcome period (pink). Water drop indicates reward delivery.

To perform 2-photon activity imaging from ensembles of DA neurons during this task, we implanted a gradient index (GRIN) lens above the VTA ^21^^,^^22^. Selective expression of GCaMP in DA neurons was achieved either by injecting a Cre-dependent AAV2/5 virus expressing GCaMP in the VTA of DAT::Cre mice, or by crossing a GCaMP reporter line with DAT::Cre mice (Fig. 1d; Supplemental Video 1 for sample imaging video; also see Extended Data Fig. 2 for relationship between spikes and fluorescence in DA neurons). In either case, an AAV2/9 mCherry virus was injected into the VTA to facilitate accurate motion correction (Extended Data Fig. 3, see Methods). Using this approach, we recorded activity of ~10-30 DA neurons simultaneously in each of 20 mice during performance of the VR task (Fig. 1e,f; n=303 DA neurons from 20 mice).

**Figure 2.**
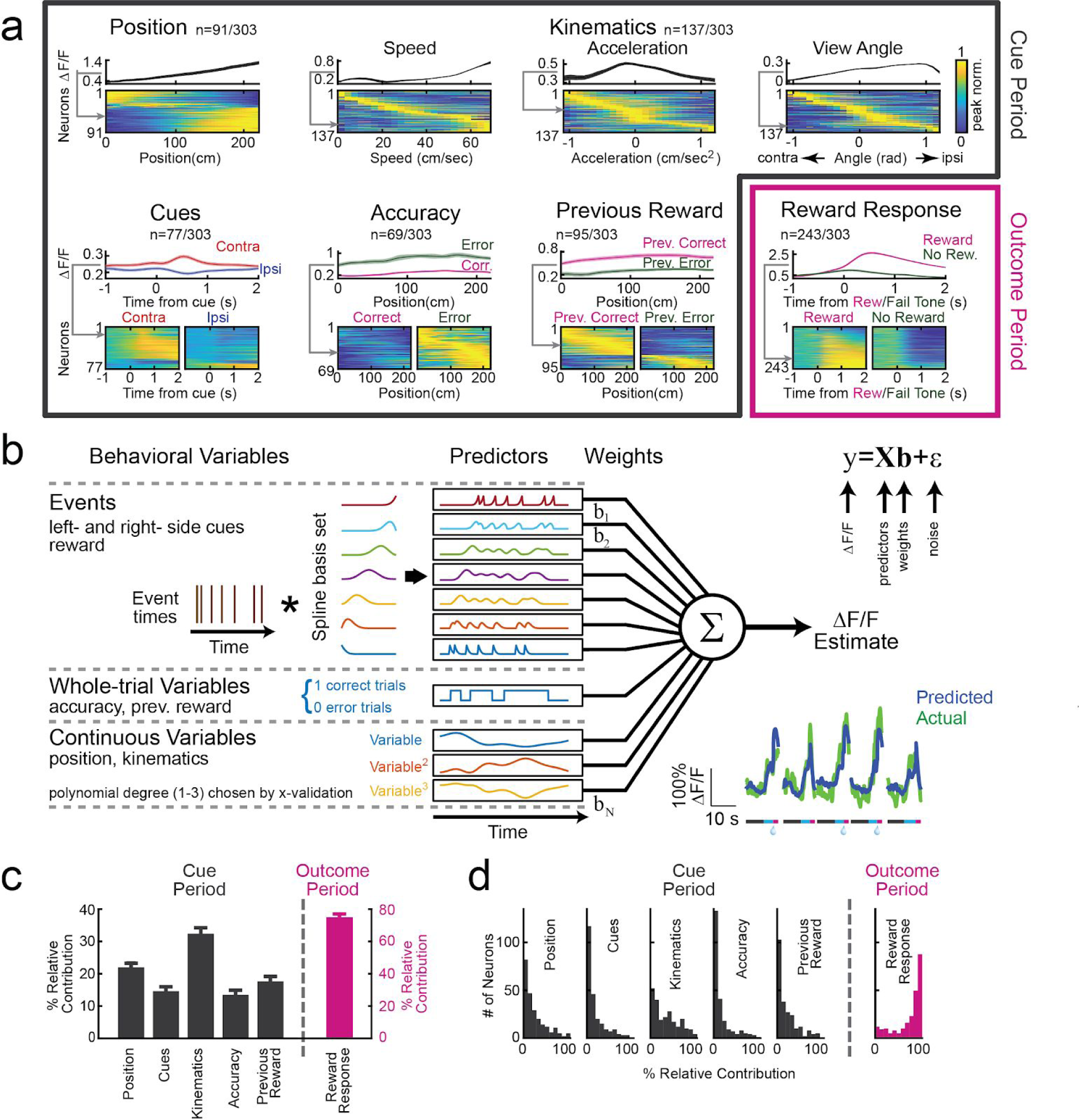
Quantifying VTA DA neuron responses to specific behavioral variables in the task. **a,** Neural activity in relation to the following behavioral variables: position along the central stem of the maze, kinematics (speed, acceleration, view angle), cues (contralateral or ipsilateral relative to the recording side), accuracy (whether or not the mouse made the correct choice at the end of the maze), previous trial reward (whether or not the previous trial was rewarded), and reward (versus no reward). The first 5 variables are quantified during the cue period while the final variable (reward) is quantified during the outcome period. For each variable, the upper panel is the average ΔF/F of an example neuron while the lower panel contains all significant neurons, with each row representing the average response of each neuron each (neuron is normalized by peak activity; grey arrow indicates the example neuron in heatmap). Statistical significance is assessed by comparing the F-statistic obtained from a nested model comparison with or without each behavioral variable to a distribution of the same F-statistic obtained from shuffled data (see Methods). In the case of position, accuracy and previous reward, the averaging is over trials. In the case of kinematics, the averaging is over timepoints. In the case of cues and reward, the averaging was across event occurrences. For the event variables (cues and reward), the average baseline activity was subtracted (in the second preceding the event). For the example neurons, false colors are s.e.m. **b,** Schematic of the encoding model used to quantify the relationship between all behavioral variables and the activity of each neuron (see Methods). Inset: predicted and actual ΔF/F across 5 trials for one neuron; more examples in Extended Data Fig. 4. **c,** Relative contribution of each behavioral variable to explained variance of the neural activity, averaged across neurons (error bars are s.e.m.). **d,** Same as **c**, but the full distribution (rather than average).

Responses of 286 out of 303 DA neurons were significantly modulated by one or more of the following variables (Fig. 2a): spatial position (n=91, 30%), kinematics (n=137, 45%), reward-predicting cues (n=77, 25%), choice accuracy (whether or not the trial resulted in reward; n=69, 23%), reward history (whether the previous trial was rewarded; n=95, 31%), and reward (n=243, 80%; significance was assessed based on nested comparisons of the encoding model described below, see Methods). The first five variables were quantified during the cue period, and the final variable (reward) was quantified during the outcome period.

During the cue period, individual neurons exhibited diverse responses to most of these variables (Fig 2a). For example, neurons that were modulated by spatial position most often exhibited upward ramps, although some displayed downward ramps. This extends, to the level of DA cell bodies, ramps previously identified with fast-scan cyclic voltammetry in the striatum ^5^^,^^23^^,^^24^ (example single trials in Extended Data Fig. 1b). Neurons that were selective to kinematics were tuned to a range of velocities, acceleration or view angles. Neurons that responded to reward-predicting cues often, but not always, displayed stronger responses to contralateral versus ipsilateral cues ^25^. Neurons that were modulated by accuracy universally displayed higher activity to error (as opposed to correct) trials, while neurons that were modulated by previous trial outcome were modulated in either direction.

In contrast to the diverse responses to many of the variables during the cue period (e.g. upward versus downward spatial ramps), most neurons responded consistently during the outcome period, with stronger responses to reward than lack of reward (Fig 2a).

Thus, for the first time, we have access to many of the behavioral variables that are thought to be relevant to DA neurons within a single behavioral paradigm. This puts us in a position to achieve our goal of understanding how the responses to these variables are organized across the DA population. To do this, we need a method to accurately quantify how much of the variance of the neural responses can be attributed to each behavioral variable individually, despite the presence of multiple behavioral variables. Towards this end, we predicted the GCaMP signal for each neuron based on all of the behavioral variables (Fig 2b; see Methods).

Briefly, to quantitatively predict GCaMP based on behavioral variables, we employed an encoding model. To derive the predictors for the model, each variable was considered either as a discrete “event” variable, a “whole-trial” variable, or a “continuous” variable. In the case of “event” variables (left cues, right cues, reward), the predictors were generated by convolving the event’s time series with a spline basis set, in order to allow flexibility in the temporal influence of cues on GCaMP. In the case of “whole-trial” variables (previous reward, accuracy), the value of the binary predictor throughout the trial indicated reward on the previous (or current) trial. In the case of “continuous” variables (position, kinematics [velocity / acceleration / view angle]), predictors included the variables raised to the first, second and third power, in order to enable flexibility in the relationship between the variable and GCaMP. This model was chosen to include behavioral variables that significantly improved predictions of neural activity, after comparing several models (see model comparisons in Extended Data Fig. 5).

Using this encoding model, we quantified the relative contribution of each behavioral variable to the response of each neuron by determining how much the explained variance declined when that variable was removed from the model (see Methods; relative contributions for example neurons in Extended Data Fig. 6). Averaged across the population, the highest relative contribution during the cue period was attributed to kinematics (32.4±1.9% of the total variance explained during the cue period), followed in descending order by spatial position (22±1.7%), previous reward (17.7±1.5%), cues (14.6±1.4%), and accuracy (13.5±1.5%, Fig. 2c,d). During the outcome period, reward contributed strongly to the response (75±2%), consistent with the large number of neurons that responded to reward (Fig 2a).

How is the relative contribution of these behavioral variables to neural responses distributed across the population? During the cue period, most behavioral variables had a small contribution to the response of each neuron, while a small subset had a large contribution. In contrast, during the outcome period, reward contributed to a large fraction of the response of most neurons (Fig. 2d). This raises the possibility that during the cue period, subsets of DA neurons are specialized to encode specific behavioral variables, while during the outcome period, most DA neurons encode reward.

To more systematically examine this idea, we performed clustering of the neurons based on the relative contributions of each behavioral variable to each neuron, using a Gaussian Mixture Model (GMM; Fig. 3a; see Methods). We found that 5 clusters of neurons gave the best (lowest) Bayesian Information Criterion (BIC) score for this data (Fig. 3a; see Methods for details on BIC score calculation). To determine if these 5 clusters in fact explained the data better than expected by chance, we compared the likelihood of the data given the clustering model to that of shuffled data, and found that the likelihood of the real data was indeed significantly higher (p<0.0001, both for null distributions generated by shuffling across behavioral variables, as well as by shuffling across neurons; Fig. 3b). Thus, we can conclude that VTA DA neurons display a statistically significant degree of functional clustering.

**Figure 3.**
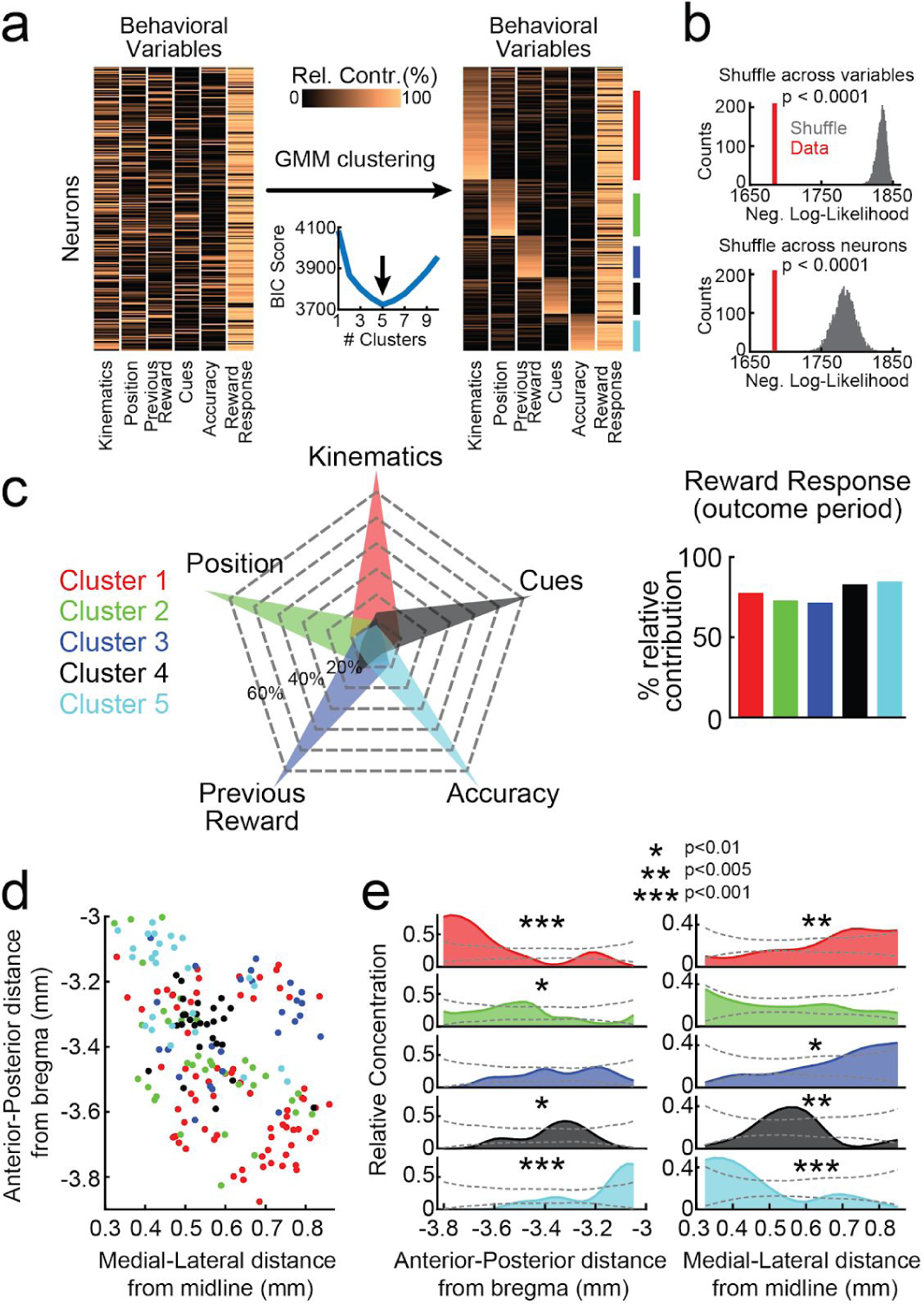
Functional and spatial organization of VTA DA neurons. **a,** The clustering procedure. Left: Relative contribution of each behavioral variable to explained variance of neural activity for each neuron before clustering (all neurons and variables are shown). Right: Same data grouped based on GMM clustering (ordered within each cluster by each neuron’s probability to belong to the cluster). The colored bars to the right of the panel denote the cluster identity. Neurons with <75% probability to belong to any cluster not assigned to a cluster (<18% of neurons unassigned). Bottom middle: BIC scores used to select the optimal number of clusters. **b,** Distributions of negative log-likelihood of cluster model for shuffled data (gray) and real data (red) indicates a significant fit of the clustering model. Top: Shuffling of relative contributions is across variables. Bottom: Shuffling is across neurons. **c,** Left: Average relative contributions of cue period behavioral variables to neural activity for each cluster. Right: average relative contribution of reward for each cluster. **d,** Recovered locations within the VTA of each neuron along the A/P and M/L axes. Cluster identity denoted by color. **e,** Relative concentration of neurons belonging to each cluster across the A/P (left) and M/L (right) axes. Dashed lines indicate the 95% confidence interval (see Methods). Relative concentration derived by normalizing the concentration of neurons belonging to each cluster by the concentration of all imaged DA neurons (see Methods). Significant spatial structure for each cluster along each axis was assessed by comparing the standard deviation of the relative concentrations of the data with that obtained from shuffled distributions (shuffling based on randomizing locations of neurons relative to cluster identity, see Methods for details; p-values are Holm-Bonferroni corrected for the 10 conditions).

Each cluster was composed of DA neurons that responded most strongly to a specific behavioral variable during the cue period. Note that this specialization does not mean that DA neurons only encoded a single variable during the cue period; in fact, many neurons also significantly encoded a 2^nd^ variable, but not as strongly (Extended Data Fig. 7). In contrast to the specialization during the cue period, all clusters were composed of neurons that had substantial reward responses (Fig. 3c). Thus, this clustering analysis provided further evidence that VTA DA neurons are specialized during the cue period, while they share a response to reward during the outcome period. Supporting the robustness of these clusters, similar clusters were obtained when the procedure was implemented independently on random halves of the trials of each neuron, and most importantly, neurons tended to be assigned to the same cluster based on each half of the trials (Extended Data Fig. 8).

We next sought to determine if the functional clusters of DA neurons were anatomically organized within the VTA. The location of each neuron was estimated based on combining histological reconstruction of the lens tract with the position of the neuron within the imaging field ^26^ (Extended Fig. 9). We observed significant dependence of cluster identity on A/P location for 4 out of the 5 clusters, and on M/L location for 4 out of the 5 clusters (Fig. 3d-e; p<0.01, comparing STD of the relative concentration of neurons within a cluster to a shuffled distribution obtained by randomly permuting the A/P or M/L location of all neurons relative to cluster identity, Holm-Bonferroni correction; see Methods). Specifically, neurons belonging to the cluster associated with kinematics were located more laterally and posteriorly (cluster 1), those associated with accuracy responses were located more medially and anteriorly (cluster 5), and neurons associated with previous reward were located more laterally (cluster 3). Thus, we find evidence for a rough anatomical map in the clusters of VTA DA neurons.

Directly correlating the A/P and M/L location of the neurons with the relative contributions of each behavioral variable led to similar findings (Extended Data Fig. 10). To ascertain that this anatomical organization cannot be explained by differences between individual mice rather than by a true dependence on location, we considered a multinomial mixed effect regression using the cluster identity of the neurons as the dependent variable, the A/P and M/L locations as fixed effects, and mouse identity as a random effect. This confirmed that anatomical location significantly predicted cluster identity (p < 0.0005, Wald test on the set of null hypotheses that all A/P coefficients in the model are equal to each other and all M/L coefficients are equal to each other).

Thus far, we have described spatial organization in the DA system based on the relative contribution of behavioral variables in explaining neural activity. A complementary approach is to examine the spatial organization of pairwise correlation between neurons. This allows us to separately consider the spatial organization of the “signal” correlation (i.e. correlations that can be explained by responses to behavioral variables; conceptually related to functional clustering in Fig. 3), and also of the “noise” correlation (i.e. neural correlations that cannot be explained by the behavioral variables). DA neurons are thought to have high noise correlations ^27^^–^^29^, but the spatial organization of these correlations has not been described.

To first confirm that DA neurons in our experiment indeed have high noise correlations, we added an additional predictor to the encoding model from Fig. 2b: a “network” predictor that reflects the activity of other simultaneously imaged neurons (for each neuron, the new predictor was the 1^st^ PCA of the ΔF/F from all other simultaneously recorded neurons; Fig. 4a). Consistent with DA neurons having high noise correlations, the performance of this new model explained a substantially higher variance of neural activity than the original model (*R^2^* from behavioral + “network” model: 50.8±1%; behavior-only model: 25.7±0.9%; Fig 4b).

**Figure 4.**
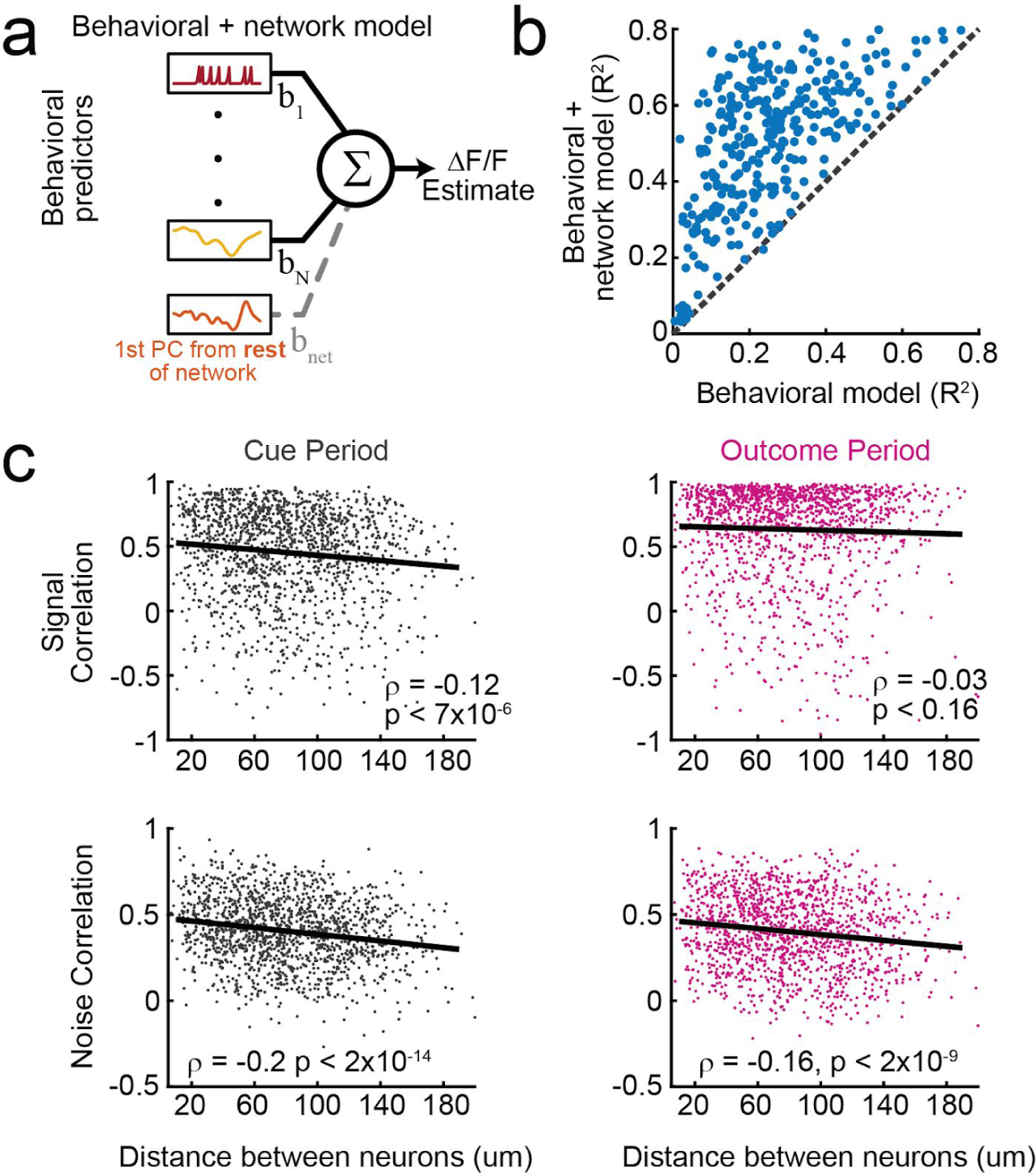
Spatial organization of signal and noise correlations in VTA DA neuron pairs. **a,** Schematic of the expanded encoding model (behavioral + network model) which includes one additional predictor compared to the model in **2b**: the 1^st^ principal component of the activity of all simultaneously recorded neurons other than the neuron being modeled. **b,** Comparison of the performance of the behavioral-only and the behavioral + network encoding models indicates high noise correlations. **c,** Signal and noise correlations for all simultaneously recorded pairs during the cue period (left) and the outcome period (right) as a function of the distance between the neurons (*n*=1492). Signal correlations decline with distance during the cue period, but not the outcome period, whereas noise correlations decline during both epochs.

We examined the spatial structure of the signal and noise correlations by considering all simultaneously recorded pairs of neurons (*n*=1492; Fig. 4c). Signal correlation was defined as the pairwise correlation between the prediction of the behavior-only encoding model for each neuron; noise correlation was defined as the pairwise correlation between the residuals of the same model. The signal correlation decreased with distance between neurons during the cue period (*r*=-0.12, *p*<7×10^-6^), but not the outcome period (*r*=-0.03, *p*<0.16). This is consistent with the results from the previous analyses, which had suggested specialized and spatially organized responses during the cue period (Fig. 3e) in contrast to widespread reward responses during the outcome period (Fig. 2a,c). On the other hand, the noise correlations decreased similarly with distance during both the cue period (*r*=-0.2, *p*<1×10^-14^) and the outcome period (*r*=-0.16, *p*<2×10^-9^), consistent with the idea that noise correlations arise from electrical synapses or shared inputs between neighboring neurons. These findings were confirmed using an alternative method for calculating noise correlations ^30^^,^^31^ (Extended Data Fig. 11).

Are the widespread reward responses in VTA DA neurons during the outcome period consistent with reward prediction error (RPE)? We first confirmed that we can replicate classic RPE during pavlovian conditioning with 2-photon imaging (Fig. 5a-c). We then sought to determine to what extent reward expectation modulates reward responses in our decision making task. In this regard, a strength of our task is that it engages two separable dimensions of reward expectation: previous trial outcome, and trial difficulty (Fig. 5d). If DA neurons reflect RPE, we would expect reward responses to be higher whenever reward expectation is low, for both dimensions of reward expectation. Indeed, across the population, reward responses were modulated by expectation in a manner that was consistent with RPE (Fig. 5e,f; median d’ = 0.1 comparing reward responses across both previous trial outcomes, p< 4×10^-11^; median d’= 0.095 comparing reward responses based on median splitting trial difficulty, p<3×10^-5^). Interestingly, across neurons, the extent of modulation by each dimension of reward expectation was (weakly) correlated, suggesting that neurons are modulated similarly by each type of RPE, rather than being specialized for one type (Fig. 5g;,*q* = .18, p < 0.005, pearson correlation between the RPE d’ values for previous trial outcome and trial difficulty for all reward responsive neurons, *n*=243). In addition, reward responses in all but one functionally defined clusters are significantly modulated by RPE (Fig. 5h,i).

**Figure 5.**
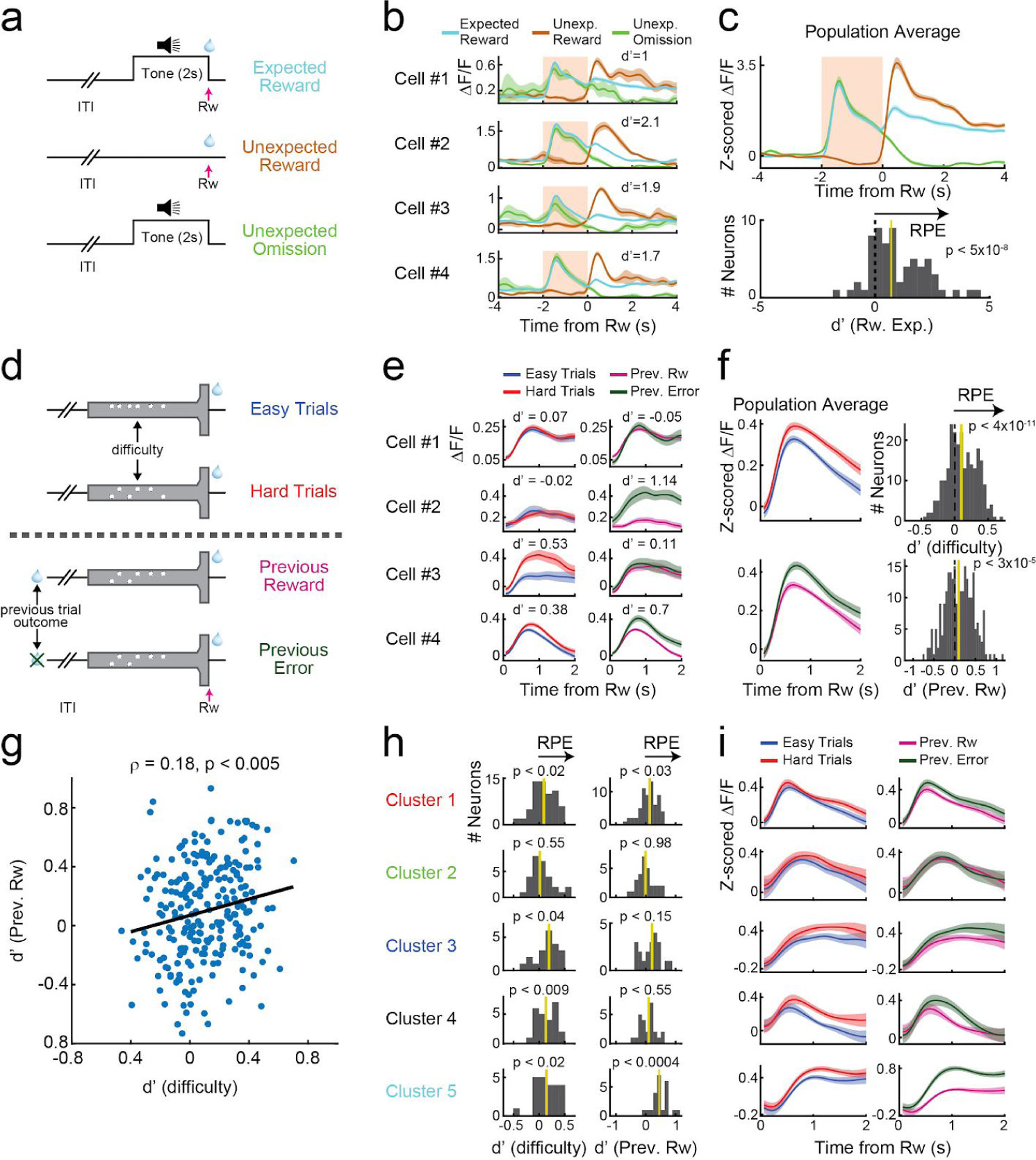
Two separable dimensions of reward expectation modulate reward responses in DA neurons during decision making. a, A Pavlovian conditioning paradigm was employed to confirm that 2-photon imaging can replicate classic RPE responses. A 2s tone preceded reward delivery during training. During the test session, in a subset trials, the tone was omitted (“unexpected reward”) or the reward was omitted (“unexpected omission”). **b,** In example cells, reward responses are modulated by expectation, consistent with RPE. d’ quantifies the RPE modulation of reward responses, and were calculated based on the average activity in the 2s following reward delivery. **c,**. Similar RPE modulation is evident in the population average (top), and in a histogram of d’ comparisons of unexpected and expected reward responses (bottom). n=8 mice and n=65neurons. **d,**. In our decision making task in VR, there are two separable dimensions of reward expectation that could theoretically modulate reward responses: trial difficulty, and previous trial outcome. **e,** Individual DA neurons can be modulated by one, neither, or both RPE dimensions. **f,** Across the population, reward responses are modulated by reward expectation in a manner that is consistent with RPE, for both dimensions of reward expectation. **g,** Across the population, there is a significant (but noisy) correlation between the 2 dimensions of RPE. **g,** Reward responses in most functionally defined clusters are significantly modulated by RPE across at least 1 dimension.

In summary, we have described organizational principles of the DA system: neurons display specialized and anatomically organized responses to non-reward variables, while the same neurons convey a less specialized reward response. These conclusions depended on combining, for the first time, a high-dimensional behavioral task (6 quantified behavioral variables) with high-dimensional neural recordings (>300 identified VTA DA neurons).

Considering the functional and anatomical organization reported here, alongside the established topography between DA neurons and their downstream targets ^25^^,^^32^^–^^34^, we can predict that specific downstream targets are likely to receive information from DA neurons about reward and only a subset of non-reward variables. Thus, this organizational structure may greatly simplify the question of how downstream circuits correctly interpret the wide range of non-reward signals encoded by midbrain DA neurons. A major open question is how downstream targets utilize these specialized non-reward signals. One possibility is that these signals reinforce downstream activity patterns related to the encoded variable, altering the probability that the behavior is repeated (in analogy to the established reinforcement function of reward responses ^35^^,^^17^^,^^36^^,^^37^). Alternatively, or in addition, they may serve to enhance ongoing activity patterns ^38^, influencing the vigor of the ongoing behavior ^39^^,^^40^, but not necessarily the probability of it being repeated in the future. New experiments will likely be designed to address these important hypotheses.

## Acknowledgments

We thank Sam Wang, Jonathan Pillow, Daniela Witten, Lucas Pinto, Scott Bolkan, David Lee, Nofar Engelhard, Ben Deverett, Alex Song, Carlos Brody, as well as the entire BRAIN COGS team and the rest of the Witten and Tank labs for comments, advice and support on this work. We also thank Esteban Engel for reagents. This research was funded by NYSCF, Pew, McKnight, NARSAD, and Sloan Foundation grants to I.B.W., and also the following NIH grants: U19 NS104648-01, DP2 DA035149-01 and 5R01MH106689-02. I.B.W. is a New York Stem Cell Foundation—Robertson Investigator.

**Extended Data Figure 1.**
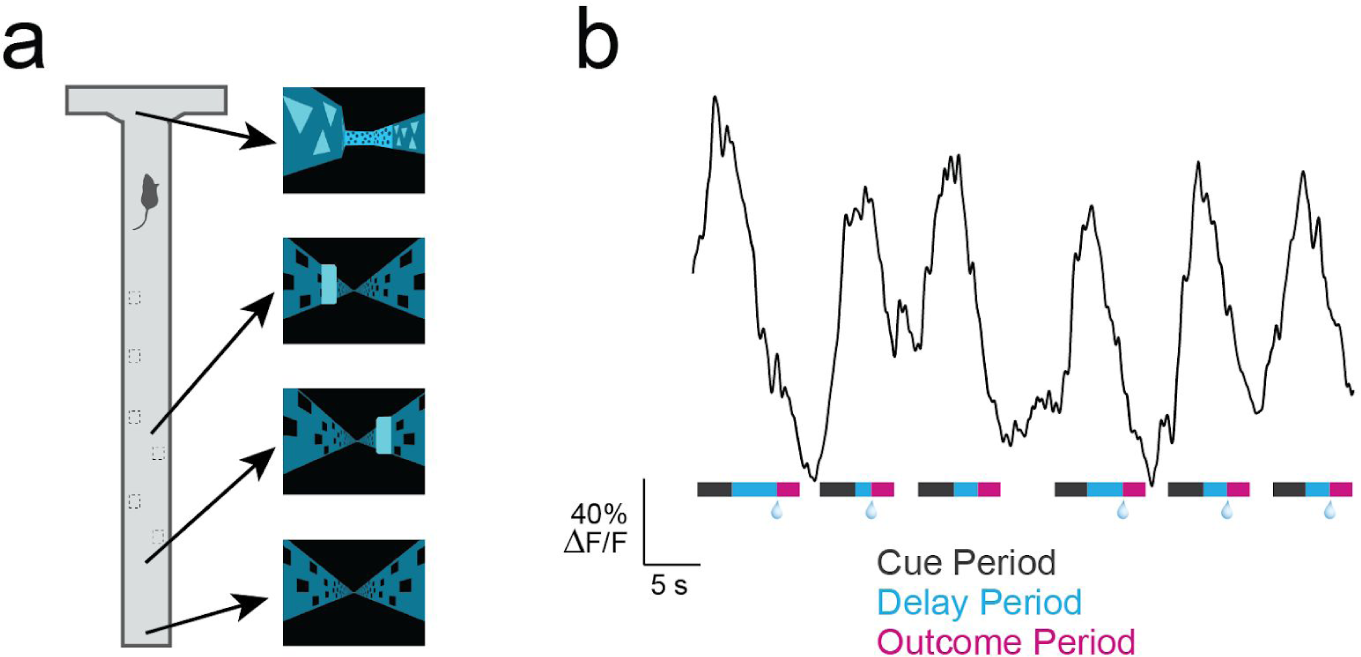
Features of the VR task. **a,** Example screenshots of the virtual world presented to the mouse in different positions along the maze. **b,** Activity trace during 6 consecutive trials of an example neuron that was significantly modulated by position in the central stem. The colored strip below the trace describes the trial epochs: cue period (gray), delay period (blue), outcome period (pink). Reward delivery is denoted by a water droplet.

**Extended Data Figure 2.**
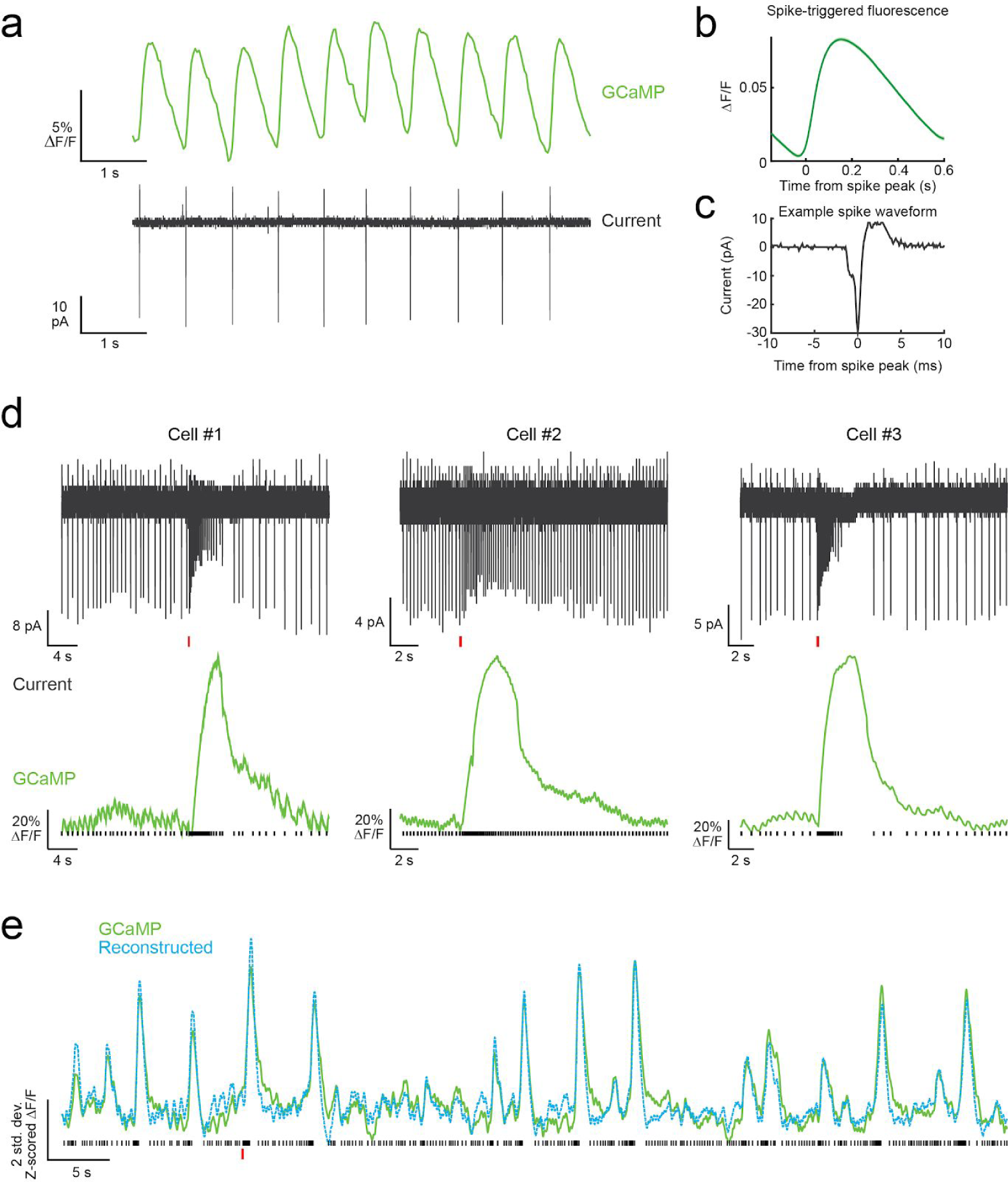
Simultaneous calcium imaging and cell-attached recording in DA neurons in the VTA of Ai148xDat::Cre mice. **a,** Relative change in fluorescence (top) and cell-attached current (bottom) recorded simultaneously. **b,** Average spike-triggered fluorescence (average over n=126 spikes). **c,** Zoomed in spike waveform for the same cell as in (a). **d,** Examples of bursts from 3 different DA cells, showing cell-attached current (top) and change in fluorescence (bottom). The spike times are shown with black bars under the fluorescence trace. The red horizontal bars under the current traces show the timing of NMDA puffs (see Methods). **e,** Example fluorescence trace (green) and reconstructed fluorescence (light blue). Fluorescence was reconstructed by convolving the spikes times (black bars, bottom) with an approximate gcamp kernel from (ref) (see Methods). Thus, this simple analysis shows that GCaMP signals in dopamine neurons can faithfully follow the spikes of DA neurons.

**Extended Data Figure 3.**
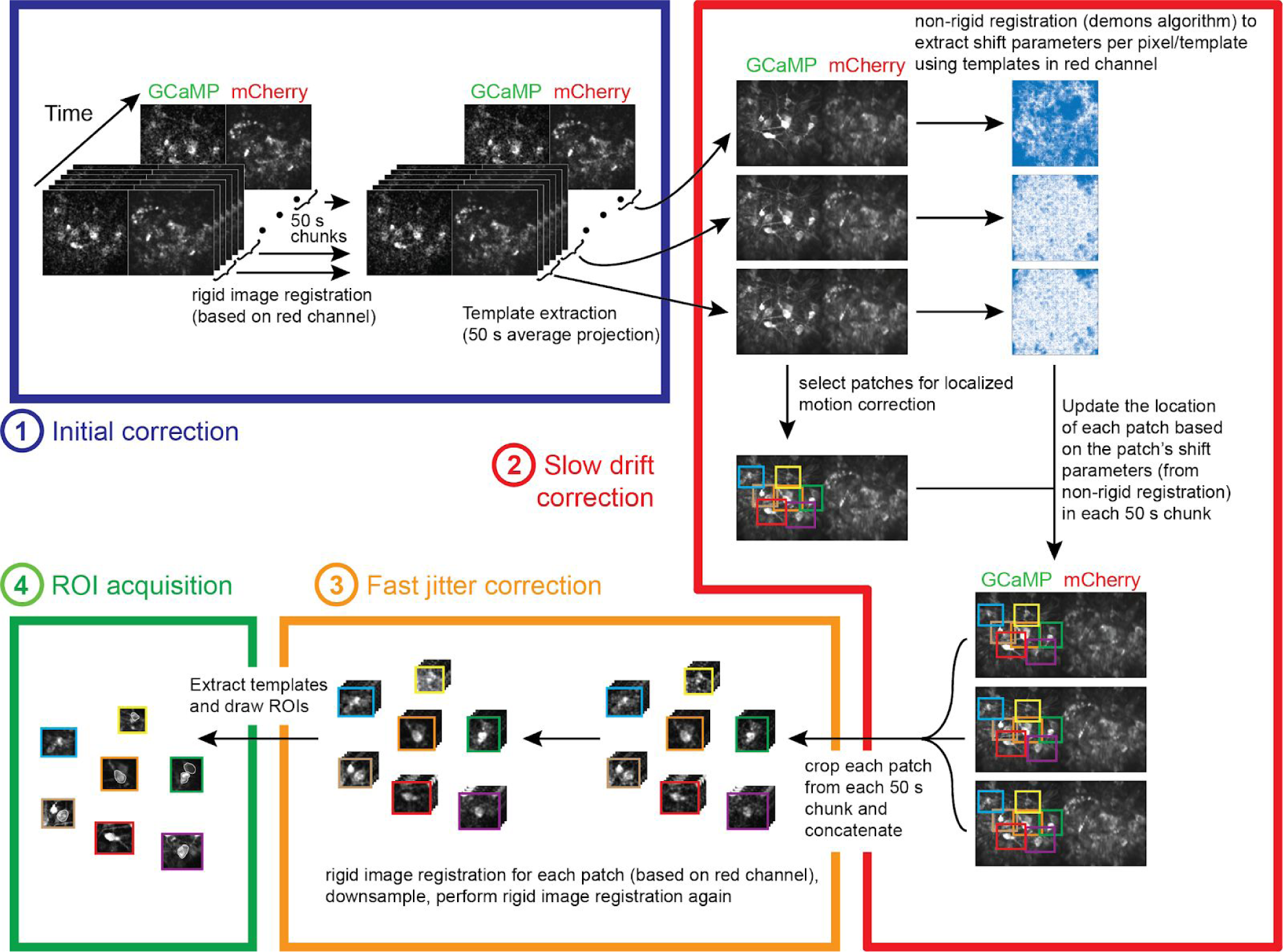
Motion correction procedure. We developed a custom motion correction procedure to compensate for both non-rigid slow drift of the field of view (timescale: 10s of min) as well as non-rigid fast motion (timescale: 10s of ms). Importantly, the procedure avoids any use of interpolation, which can produce motion artifacts. The procedure consists of the following main steps: **1** (blue box) the entire movie is divided in non-overlapping 50 s chunks; in each chunk we perform rigid motion correction using standard cross-correlation methods (on the red channel). The template for each chunk is calculated by dividing the chunk into non-overlapping sections of 100 frames, calculating the mean image of each section, and obtaining the median of the mean images. **2** (red box) we use a non-rigid algorithm for image registration to align all the templates. The algorithm outputs shift parameters for every pixel and template. Separately, we manually draw patches that include neurons of interest in the first template. For each template, we use the shift parameters of all the pixels in each patch to estimate the average motion of the patch. We use that information to crop the patch from each 50 s chunk of the movie. **3** (orange box) we perform rigid motion correction (as above) on the concatenated patch movies, down-sample by a factor of 2 (to increase the signal strength) and then perform rigid motion correction again. **4** (green box) we extract the patch templates by using the mean projection, and hand draw ROIs of the objects of interest. See Methods for a detailed explanation of motion correction algorithm, and see Supplemental Video 1 for an example video before and after correction.

**Extended Data Figure 4.**
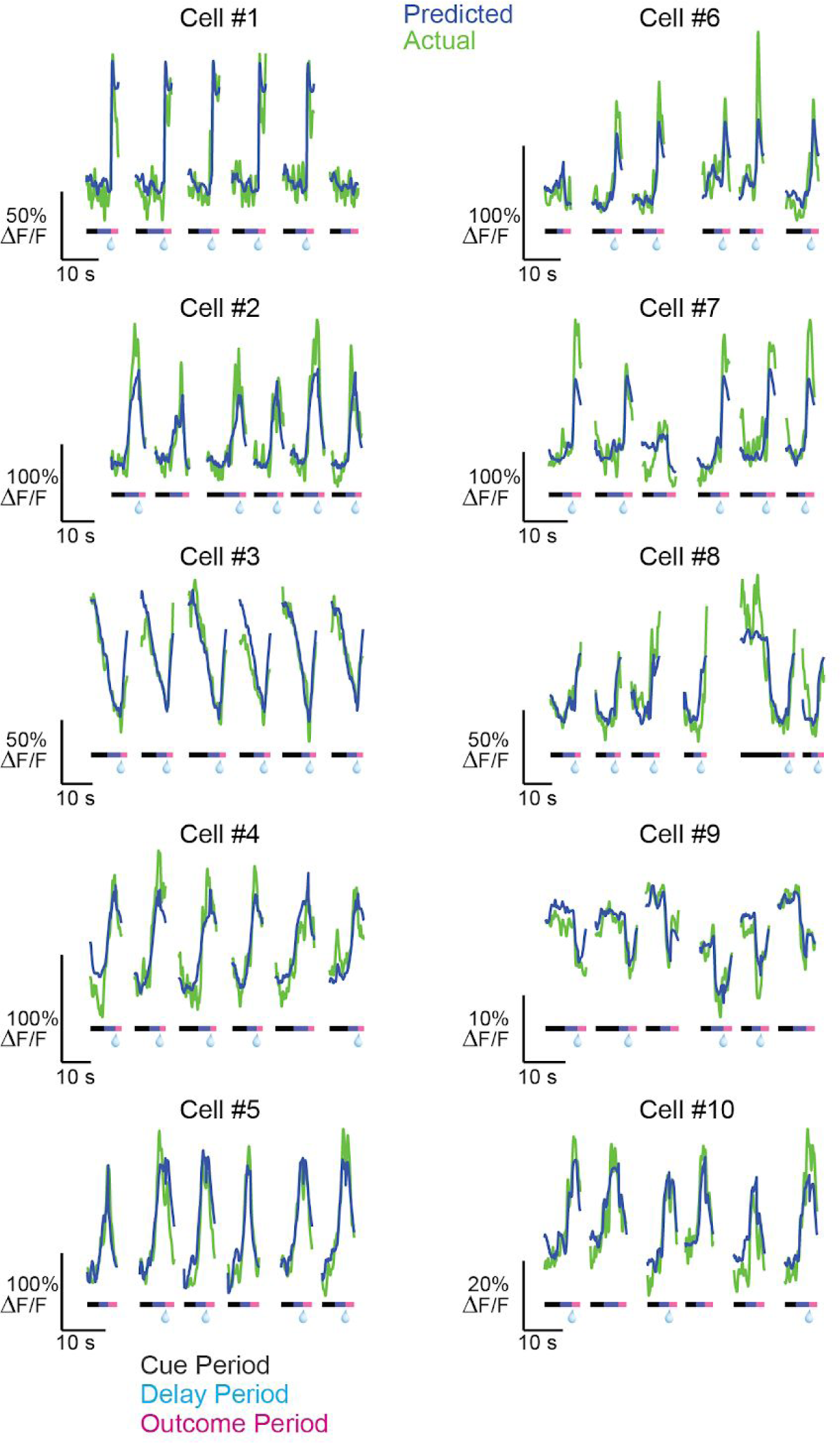
GCaMP fluorescence and corresponding predictions from example DA neurons. ΔF/F traces for 10 example neurons during 6 consecutive trials (green). Overlaid are the predictions of the behavioral model for these trials (blue). The colored strip below each trace denotes the trial epochs: cue period (gray), delay period (blue), outcome period (pink). Reward delivery is denoted by a water droplet.

**Extended Data Figure 5.**
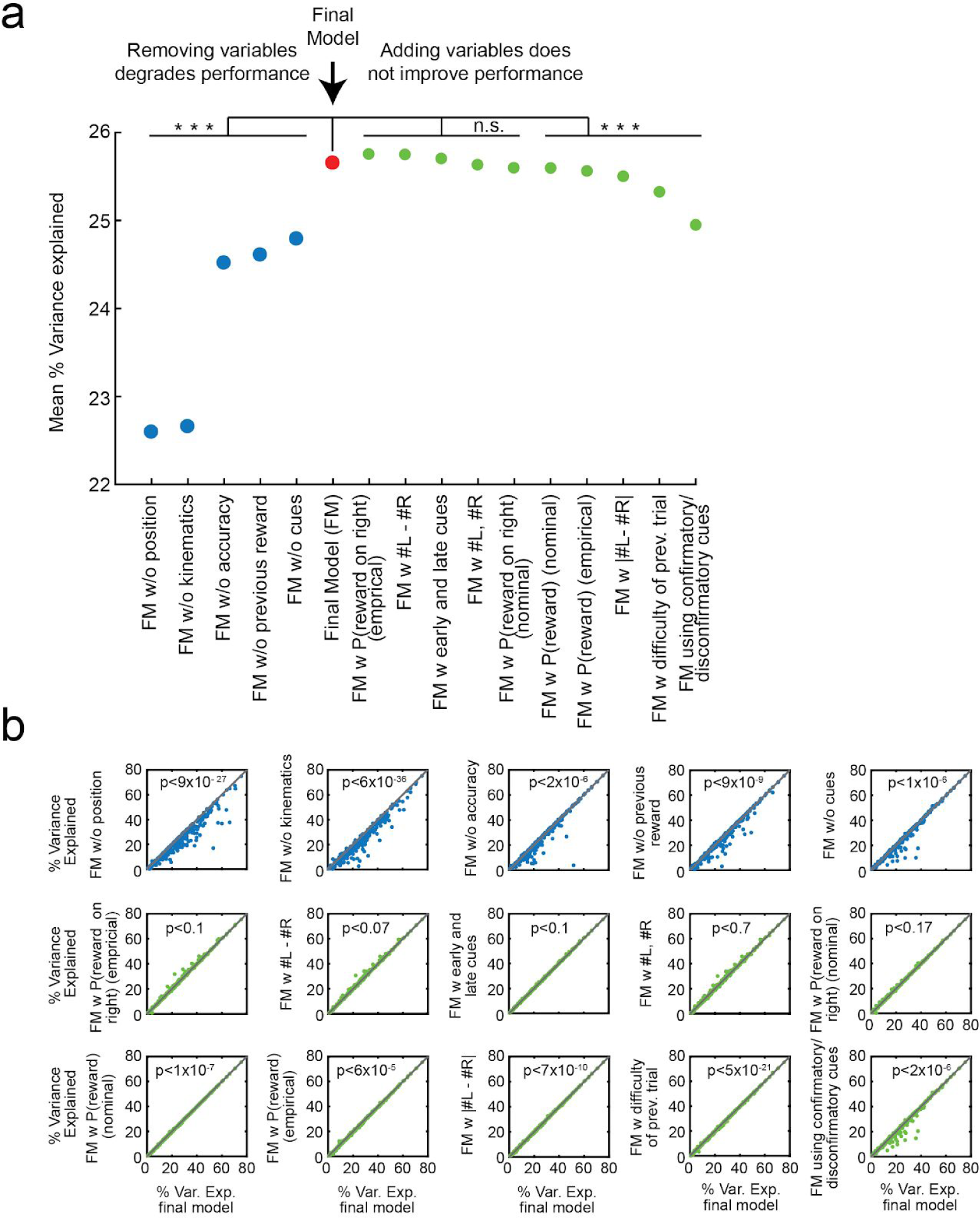
Selection of the encoding model. **a,** Mean (across neurons) of percent variance explained (tested with crossvalidation) by the final model (red) and other models where a variables was either removed (blue) or added (green). See Methods for descriptions of all variables that were tested. All models for which a variable was removed from the final model performed significantly worse, based on comparing R2 for all neurons (p<2×10^-6^, paired t-test, Holm-Bonferroni correction for all model comparisons). For models where variables were added to those in the final model, the performance either did not exhibit a significant difference, or was degraded. See Methods for complete description of all models. **b,** Comparison of performance for all neurons of the final model (x-axis) and all the other models. Each panel shows the comparison with one model; significance of the paired t-test (After Holm-Bonferroni correction) is shown in each panel.

**Extended Data Figure 6.**
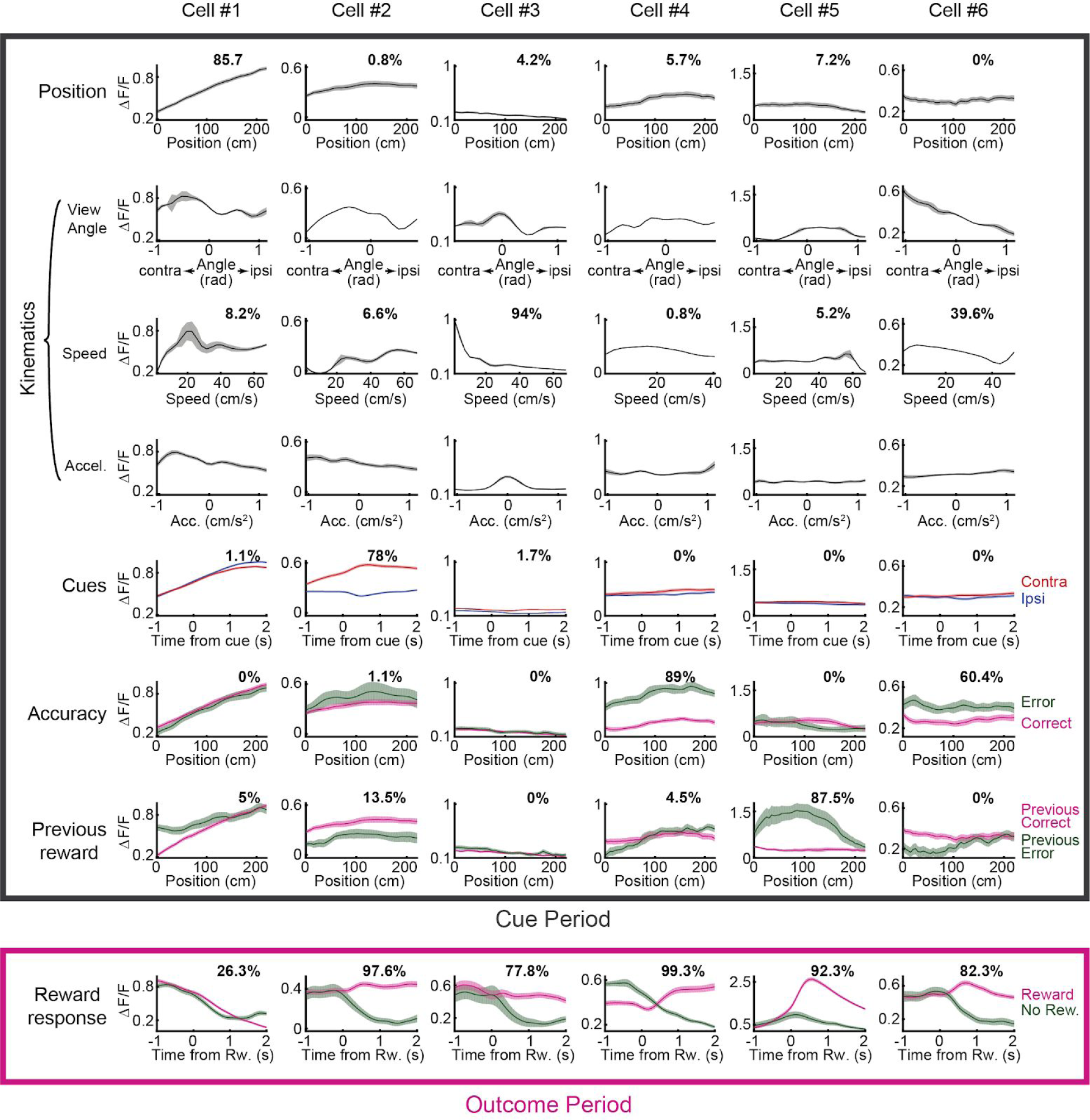
Average activity and relative contributions of different behavioral variables for several example cells. The panels show activity averages time-locked to different behavioral variables for 6 example cells. The percentage of relative contribution of the corresponding behavioral variable to the activity of each cell is displayed in each panel.

**Extended Data Figure 7.**
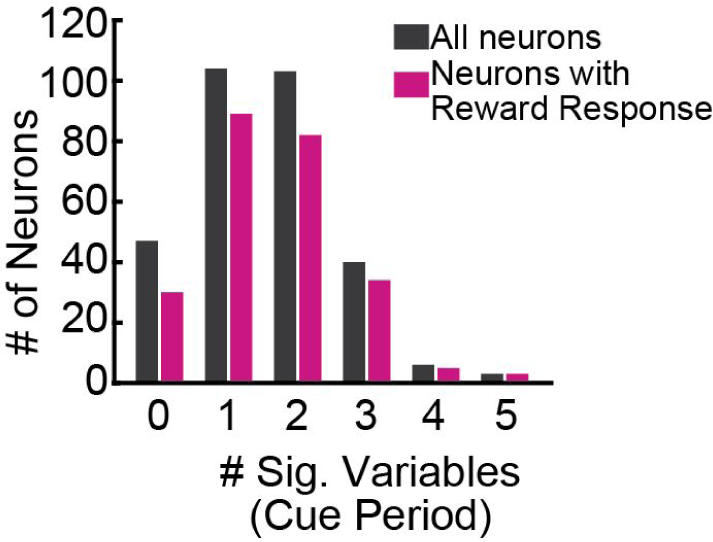
Number of behavioral variables significantly encoded per neuron across the DA population. The histogram of the number of behavioral variables during the cue period for which neurons had a significant response to is shown for all neurons (grey) and for the subset of neurons that had a significant reward response (pink). Statistical significance is assessed by comparing the F-statistic obtained from a nested model comparison with or without each behavioral variable to a distribution of the same F-statistic obtained from shuffled data (see Methods).

**Extended Data Figure 8.**
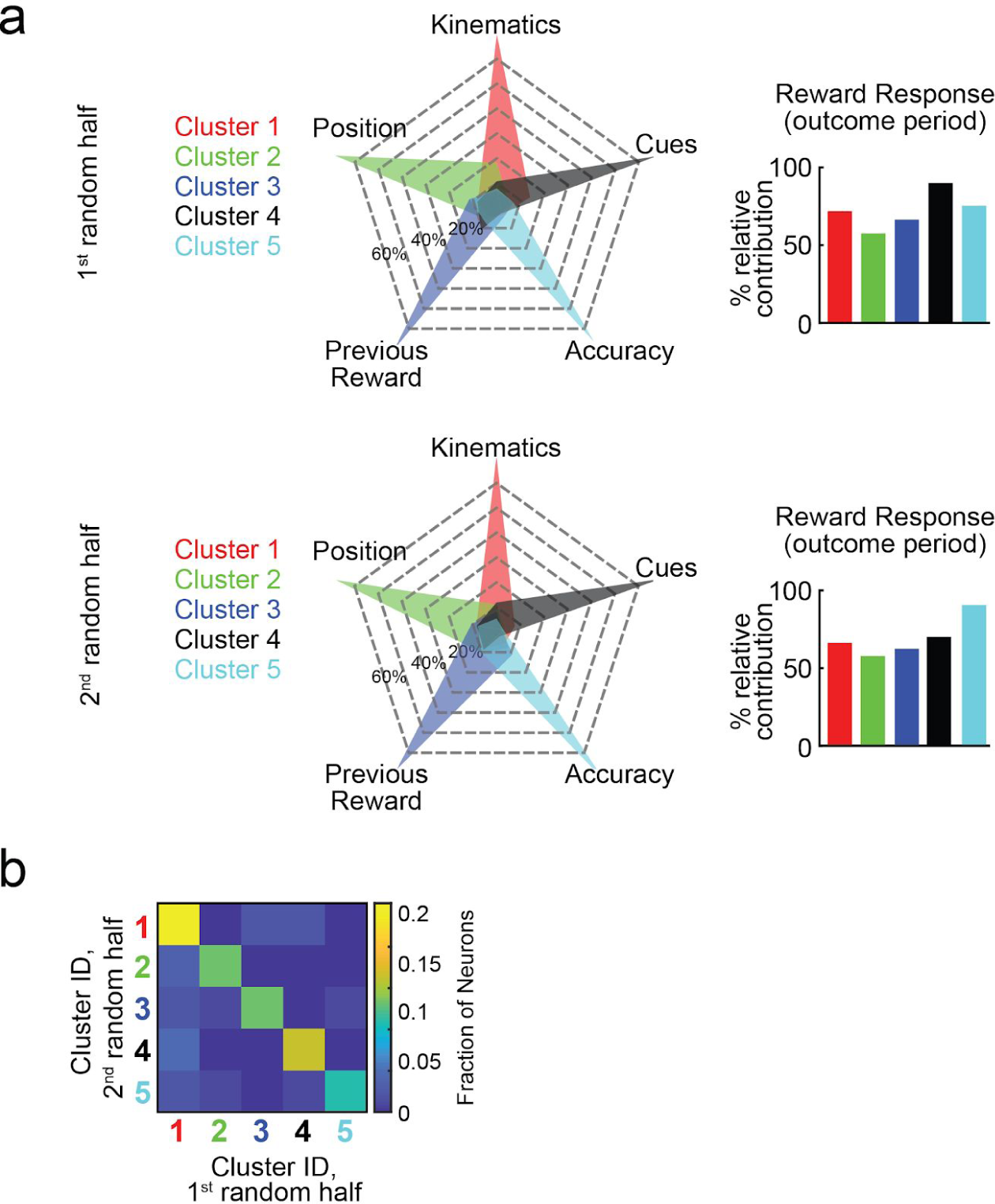
Robustness of clustering results, assessed based on comparing clustering results for each neuron between each half of trials. **a,** Average relative contributions of clusters obtained by separately analyzing two random halves of the trials for each neuron. Correlations between the average relative contributions in each cluster across the two sets are as follows: Position: *ϱ* = .99, p < 8×10^-5^. Cues: *ϱ* = .99, p < 4×10^-4^, Kinematics: *ϱ* = .99, p < 2×10^-4^. Accuracy: *ϱ* = .99, p < 3×10^-4^. Previous Reward: *ϱ* = .99, p < 0.001. Reward Response: *ϱ* = .48, p < 0.42. **b,** Normalized confusion matrix for the cluster identities of each neuron, obtained by clustering the two random halves of the data. The main diagonal represents neurons for which the cluster identities matched (79.1%). Note that chance level of matching is 20%. The matrix was calculated for neurons for which a cluster was assigned in the procedures for both halves of the data (>75% probability to belong to a cluster, n=91).

**Extended Data Figure 9.**
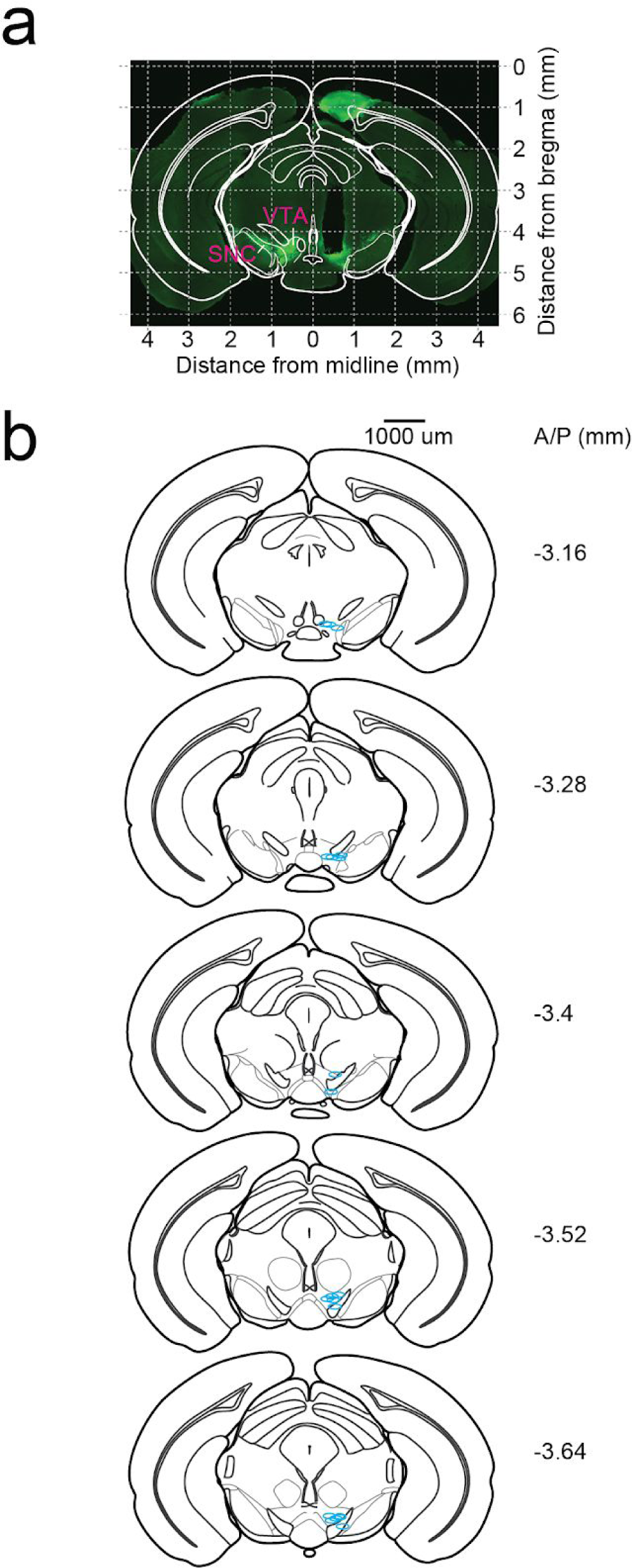
Summary of GRIN lens locations. **a,** Example of lens location recovery. Coronal histological slices stained for Tyrosine hydroxylase (green) were aligned to atlas image sections. The center of the lens was marked and its position in common coordinates was recovered by using the atlas measurements. **b,** Recovered centers of GRIN lenses from all mice (blue ellipses) are shown on top of the atlas images.

**Extended Data Figure 10.**
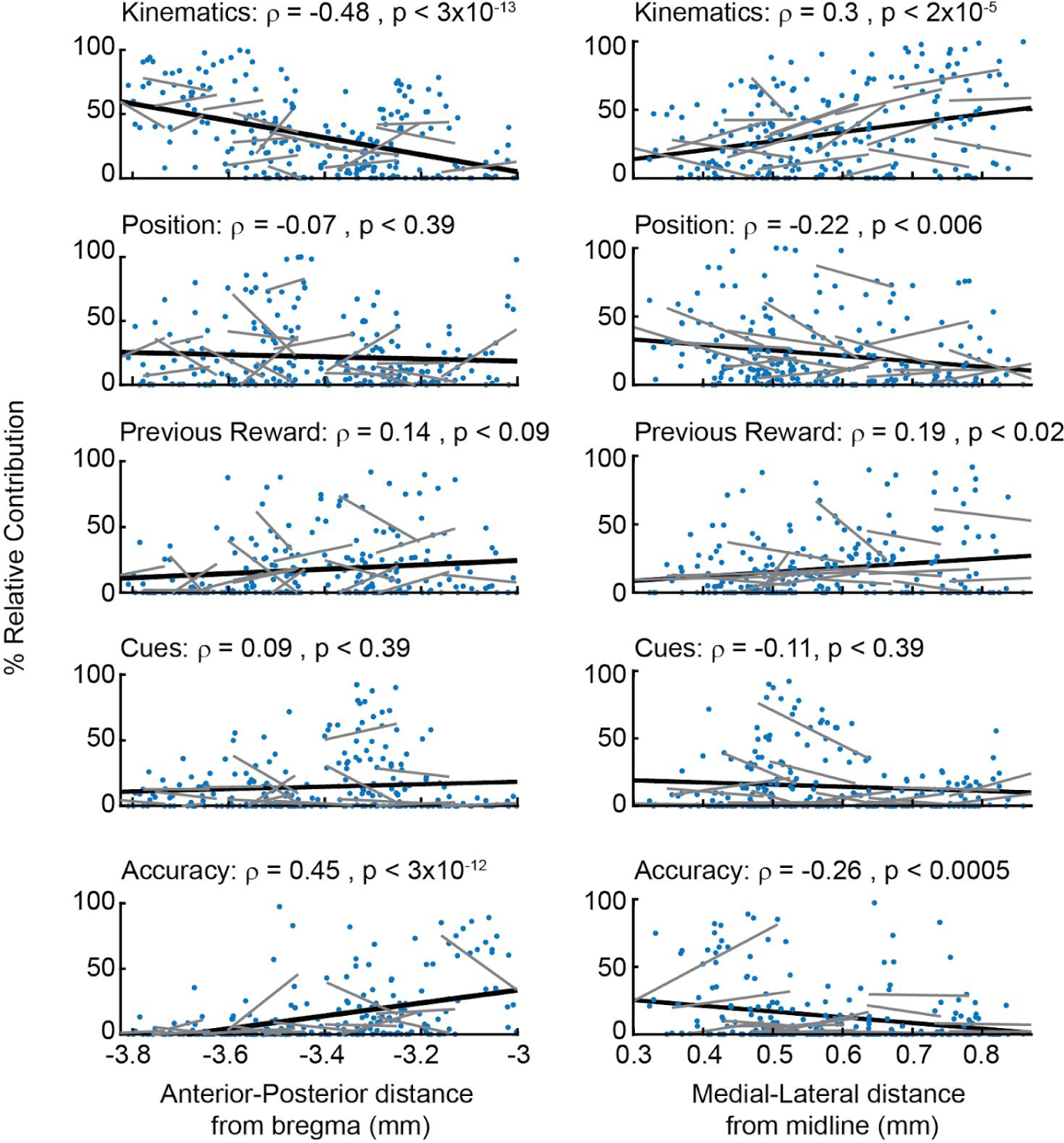
Relative contributions of each behavioral variable as a function of neuron location along the A/P and M/L axes, and mixed effect model to account for individual differences. In each row, the relative contribution of a behavioral variable is correlated with the A/P (left) or M/L (right) locations. The correlation value and significance (after Holm-Bonferroni correction for all tests) is shown in the panel. The linear fits of the entire population is shown by a black line, and linear fits of neurons belonging to individual mice (which had more than 5 neurons) are shown by gray lines. To test statistically for a dependence on position while accounting for individual differences, we additionally considered a mixed effect model, in which the relative contribution was the dependent variable, the A/P, M/L locations and their interaction were independent fixed effects, and the mouse identity was a random effect. We obtained the following p-values (F-test for the fixed effects): Kinematics: p<0.0007, Position: p<0.12, Previous Reward: p<0.007, Cues: p<0.36, Accuracy: p<0.002. This indicates that the relative contributions are dependent on spatial location in the case of kinematics, previous reward and accuracy, even when accounting for the “average offset” from individual mice. In the case of Previous Reward, but not the other variables, the mixed effect model gave better fits when including the interaction between M/L and A/P. The improved fit with the interaction explains the lower p-values for the full mixed effect model for Previous Reward compared to the direct correlation between the relative contribution of Previous Reward with A/P or M/L alone, as shown in this figure.

**Extended Data Figure 11.**
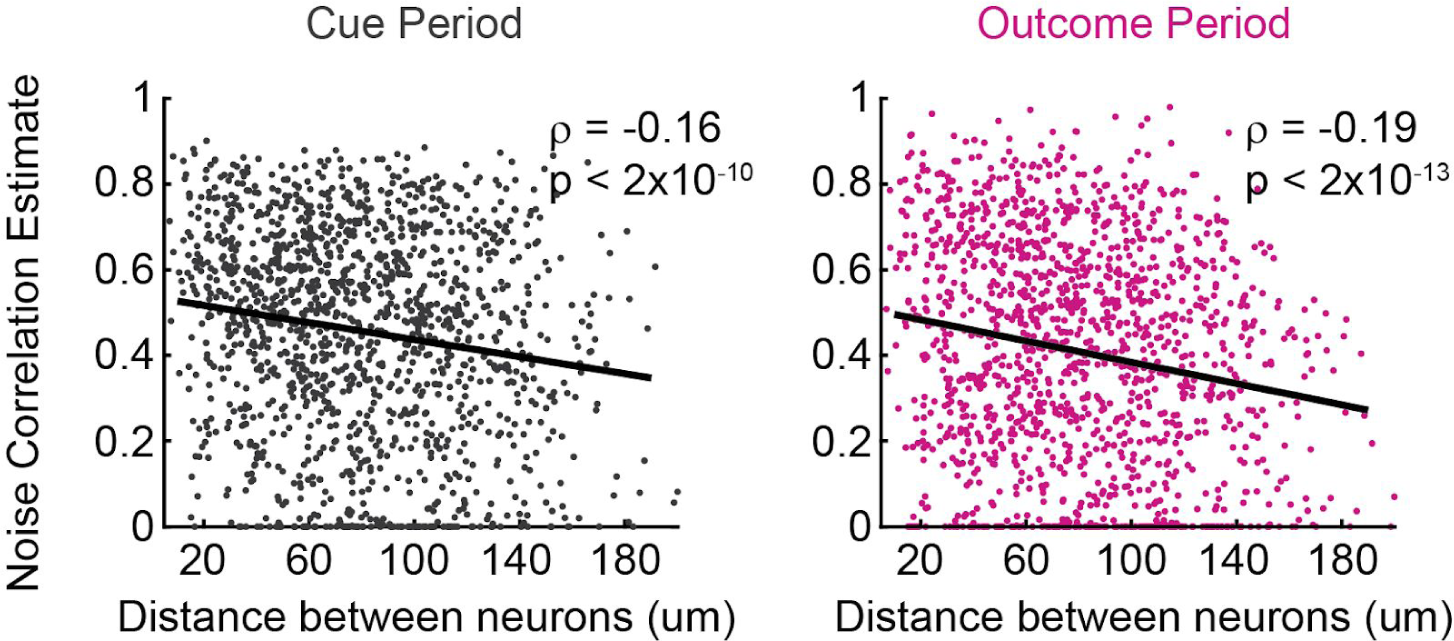
Noise correlations estimated by an alternative method. Here, noise correlations were estimated by calculating the increase in variance explained by the behavioral-only encoding model when the second neuron activity was added to it as a predictor ^30^^,^^31^ . The noise correlation estimate is shown for all neuronal pairs (n=1492) during the cue period (left) and outcome period (right).

**Extended Data Figure 12.**
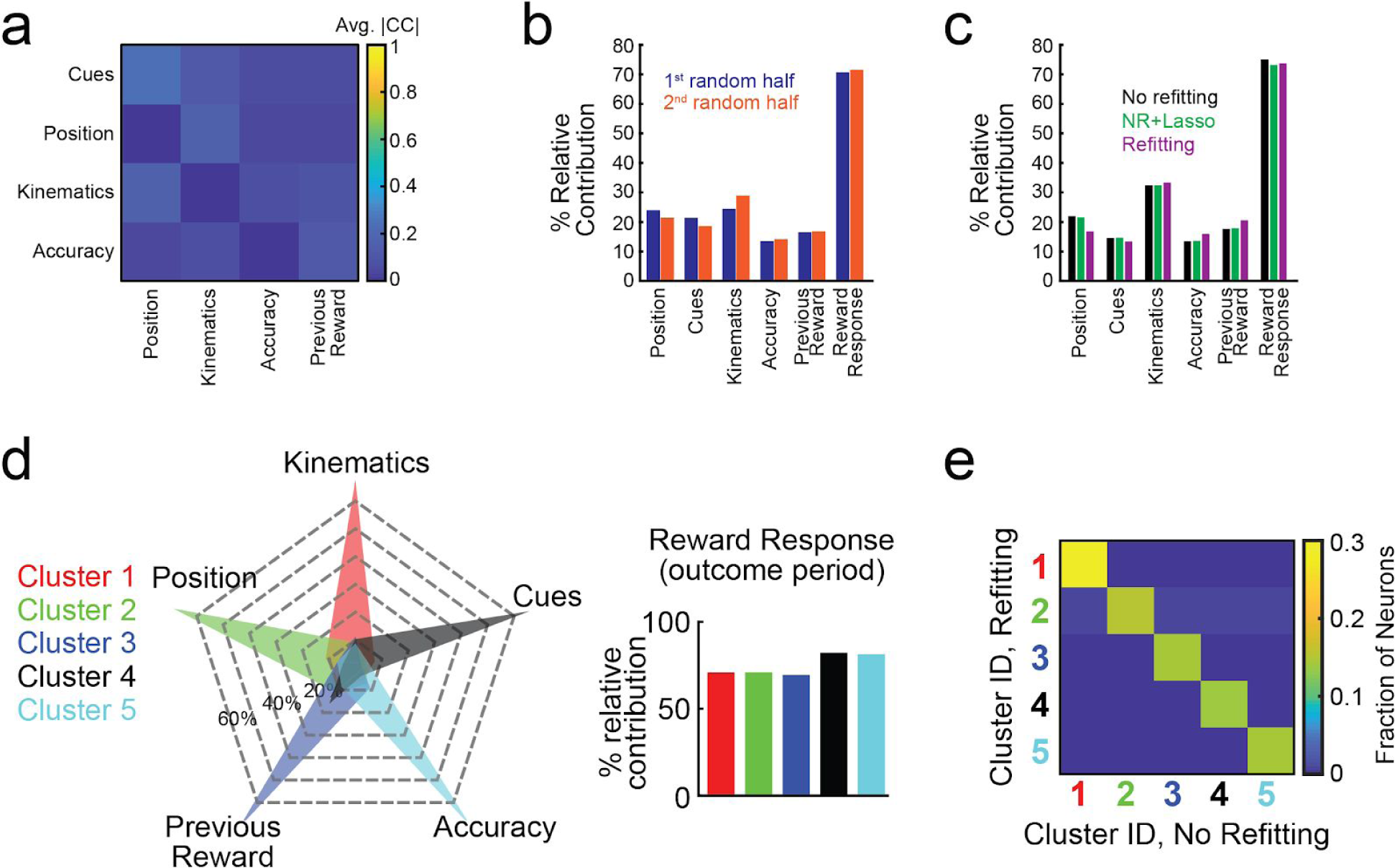
Validation of the encoding model. **a,** Average absolute value of the correlations for all pairs of predictors across all behavioral variables during the cue period (average across all predictor pairs and mice). **b,** Average relative contributions assessed separately using two random halves of the data. For each neuron we randomly divided all the trials where the neuron was recorded into 2 separate subsets while matching the number of rewarded and previously rewarded trials between the subsets. Each subset of trials was then used to calculate the relative contributions of the behavioral variables. (*ϱ* = .99, p < 3×10^-4^ for all behavioral variables, *ϱ* = .8, p < 0.11 when omitting the reward response contributions). **c,** Average relative contributions assessed separately using 3 different approaches: 1- No refitting (NR; used in the paper). 2- No refitting + LASSO regularization (NR+L). 3- Refitting (R). Correlations between the results of the different approaches are as follows: *ϱ*(NR,NR+L) = 1, p < 7×10^-9^. *ϱ*(NR,R) = .99, p < 1×10^-4^. *ϱ*(NR+L,R) = .99, p < 8×10^-5^. When omitting the reward response contributions: *ϱ*(NR,NR+L) = 1, p < 2×10^-5^. *ϱ*(NR,R) = .91, p < 0.04. *ϱ*(NR+L,R) = .92, p < 0.03. Lasso regularization was applied using the ‘lasso’ function in Matlab; the mean square error (MSE) of the model was estimated using 5-fold crossvalidation, and we chose the lambda value that minimized the MSE. The results with lasso regularization were almost identical to the result without regularization, suggesting that there was not significant overfitting in our model. **d,** Results of the clustering analysis performed on the contributions calculated using the refitting approach. Left: The average relative contributions of cue period behavioral variables to neural activity for each cluster. Right: average relative contribution of the reward response for each cluster. **e,** Normalized confusion matrix for the cluster identities of each neuron, obtained by comparing the clustering of the relative contributions based on either the No-refitting or the Refitting approach (see Methods for description of 2 approaches). The main diagonal represents neurons for which the cluster identities matched (97.8%).

**Extended Data Table 1.** Details of the shaping procedure. The tables lists the parameters of the mazes progressively used during the shaping of the behavior. The “permanent cues” field indicates if the cues were presented at the beginning of the trial; otherwise, each cue was presented when the mouse was 10 cm away from its location. High- and Low-cue-probability side poisson means indicate the means of the poisson distribution from which the number of cues presented on each side were drawn (at least 1 cue was always drawn); “none” indicates that no cues were presented for the low-probability side on any trial in that maze.

|  |  |  |  |  |  |  |  |  |  |
| --- | --- | --- | --- | --- | --- | --- | --- | --- | --- |
| Maze # | 1 | 2 | 3 | 4 | 5 | 6 | 7 | 8 | 9 |
| Start region length (cm) | 5 | 30 | 30 | 30 | 30 | 30 | 30 | 30 | 30 |
| Cue region length (cm) | 45 | 100 | 200 | 220 | 220 | 220 | 220 | 220 | 220 |
| Delay region length (cm) | 10 | 20 | 20 | 50 | 80 | 80 | 80 | 80 | 80 |
| Permanent cues (y/n) | y | y | y | y | y | n | n | n | n |
| High-cue-probability side poisson mean | 0.45 | 4 | 10 | 11 | 8.8 | 8.8 | 7.1 | 6.8 | 6.4 |
| Low-cue-probability side poisson mean | none | none | none | none | none | none | 0.6 | 0.9 | 1.3 |

**Supplementary Video 1.** Raw (left) and motion-corrected (right) video of DA neurons imaged through a GRIN lens during behavior. For visualization, the movie was sped up by a factor of 5 by taking a rolling average of 5 frames and downsampling by a factor of 5.

## Methods

### Animals and surgery

All experimental procedures were conducted in accordance with the National Institutes of Health guidelines and were reviewed by the Princeton University Institutional Animal Care and Use Committee (IACUC). A total of 28 mice were used in this study. For the virtual reality experiments, we used either male DAT::IRES-Cre mice (n=14, The Jackson Laboratory strain 006660) or male mice resulting from the cross of DAT^IRES*cre*^ mice and the GCaMP6f reporter line Ai148 mice^41^ (n=6, Ai148xDAT::cre, The Jackson Laboratory strain 030328). For the pavlovian conditioning experiments, we used male and female Ai148xDAT::cre mice (n=8). Mice were maintained on a 12-hour light on – 12-hour light off schedule. All procedures were conducted during their light off period. Mice were 2-6 months old.

Mice between 8-12 weeks underwent sterile stereotaxic surgery under isoflurane anesthesia (3-4% for induction, .75-1.5% for maintenance). The skull was exposed and the periosteum removed using a delicate bone scraper (Fine Science Tools). The edges of the skin were affixed to the skull using a small amount of Vetbond (3M). We injected 800nl of a viral combination of AAV5-CAG-FLEX-GCaMP6m-WPRE-SV40 (n=12) or AAV5-CAG-FLEX-GCaMP6f-WPRE-SV40 (n=2; U Penn Vector Core) with 1.6×10^12^/mLtiter and AAV9-CB7-CI-mCherry-WPRE-rBG (U Penn Vector Core) with 2.3×10^12^/mL titer. Two such injections were made at stereotactic coordinates: 0.5 mm lateral, 2.6 or 3.8 mm posterior, 4.7 mm in depth. After the injections, we implanted a 0.6 mm diameter GRIN lens (GLP-0673, Inscopix or NEM-060-25-10-920-S-1.5p, GrinTech) in the VTA (coordinates shown in Extended Data Fig. 9) using a 3D printed custom lens holder. After implantation, a small amount of diluted metabond cement (Parkell) was applied to affix the lens to the skull using a 1ml syringe and 18 gauge needle. After 20 minutes, the lens holder grip on the lens was loosened while the lens was observed through the microscope used for surgery to ascertain there was no movement of the lens. Then, a previously described titanium headplate was positioned over the skull using a custom tool and aligned parallel to the stereotax using an angle meter ^42^ . The headplate was then affixed to the skull using metabond. A titanium ring was then glued to the headplate using dental cement blackened with carbon.

### Virtual reality behavioral system

In order to enable a navigation-based decision making task under head-fixed conditions, we used a virtual reality (VR) system similar to that described previously ^43^^,^^44^ (Fig. 1a). Mice were held head-fixed under a two-photon microscope using two custom headplate holders and ran on an air-supported, Styrofoam spherical treadmill that was 8-inch in diameter. We found that the precise alignment of the mouse on top of the sphere was important for maintaining good behavioral performance; therefore, we used a custom alignment tool for this purpose. The sphere’s movement were measured using an optical flow sensor (ADNS3080) located underneath the sphere and controlled by an Arduino Due; this information was sent to the VR computer, running the ViRMEn software engine ^45^ (https://pni.princeton.edu/pni-software-tools/virmen) under Matlab, which displayed and controlled the VR environment. The measured sphere displacements (*dX* and *dY*, where Y is parallel to the long stem of the T-maze) resulted in translational displacements in the virtual environment of equal length in the corresponding axis. The speed of the mouse was given by 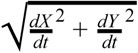, where *dt* was the time elapsed from the previous sampling of the sensor. The mouse acceleration was the moment-by-moment change in speed. The mouse view angle in the virtual world was calculated as follows: first, we calculated the current displacement angle as: ω = *atan*2(− *dX* · *sign*(*dY*), |*dY* |) . Then, the rate of change of the view angle (θ) was given by:

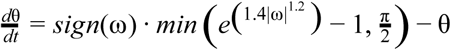

This exponential function was tuned to stabilize trajectories during the long stem of the maze, while allowing sharp turns into the maze arms (see ^20^ for more details).

The display was projected using a DLP projector (Mitsubishi HD4000) running at 85 Hz onto a custom toroidal screen with a 270° horizontal field of view. Reward delivery was accomplished by sending by a TTL pulse from the VR computer to a solenoid valve (NResearch) which released a drop of a water to a lick tube located slightly in front and below the mice’s mouth. The tone signifying trial failure was played through conventional computer speakers (Logitech). The setup was enclosed in a custom-designed cabinet built from optical rails (Thorlabs) and lined with sound-absorbing foam sheeting (McMaster-Carr).

### Optical imaging and data acquisition

Imaging was performed using a custom-built, VR-compatible two-photon microscope ^44^. The microscope was equipped with a pulsed Ti:sapphire laser (Chameleon Vision, Coherent) tuned to 920nm. The scanning unit used a 5mm Galvanometer and an 8 kHz resonant scanning mirror (Cambridge Technologies). The collected photons were split into two channels by a dichroic mirror (FF562-Di03, Semrock). The light for the green and red channels respectively were filtered using bandpass filters (FF01-520/60 and FF01-607/70, Semrock), and then detected using GaAsP photomultiplier tubes (pmts, 1077PA–40, Hamamatsu). The signal from the pmts was amplified using a high speed current amplifier (59-179, Edmund). Black rubber tubing was attached to the objective (Zeiss 20×, 0.5 NA) as a light shield covering the space from the objective to the titanium ring surrounding the GRIN lens. Double distilled water was used as the immersion medium. The microscope could be rotated along the medial-lateral axis of the mice which allowed alignment of that optical axes of the microscope objective and GRIN lens as described previously for microprism imaging ^44^. Control of the microscope and image acquisition were performed using the ScanImage software (Vidrio Technologies; ^46^) that was run on a separate (scanning) computer. Images were acquired at 30 Hz at a resolution of 512 x 512 pixels. Average beam power measured at the front of the objective was 40-60 mW. Synchronization between the behavioral logs and acquired images was achieved by sending behavioral information each time the VR environment was refreshed from the VR computer to the scanning computer via an I2C serial bus; behavioral information was then stored in the header of the image files.

### Behavioral training

Seven days after the surgery, mice were started on a water restriction protocol, with a daily allotment of water of 1 – 1.5 ml. Mice were monitored for signs of dehydration or drops in body mass below 80% of the initial value. If any of these conditions occurred, mice were given *ad libitum* access to water until recovering. The animals were handled daily from the start of water restriction. 5 days after starting water restriction and handling, mice began training in the behavioral setup. Training consisted of a shaping procedure with 9 levels of T-mazes with progressively longer stem length and cognitive difficulty (Extended Data Table 1). After shaping concluded, in each session the first few trials (5-30) were warm-up trials drawn from mazes 5-8, and then trials from the final maze (#9) were used for the remainder of the session; only the portion of the session with maze #9 trials was used for all analyses in the paper. The mice typically received their daily allotment of water during task performance; if not, the remainder was provided to them at the end of the day.

### Details of the behavioral task

At the beginning of each trial, mice were presented with the start of a virtual T-maze. After 30 cm (Start region) the cue region began, in which cues randomly appeared on either side of the corridor. The number of cues presented were sampled from a Poisson distribution, with means of 6.4 to one of the sides, and 1.3 to the other. In order to obtain better estimation of the psychometric curves, we additionally oversampled easy trials by having 5% of trials with a difference in # cues between the sides of 12 or more (using the same probability distributions). The identity of the high-cue-probability and low-cue-probability sides (left or right) were recalculated each trial to randomize the task and avoid side bias ^20^. The locations of the cues were randomly assigned along the cue region, such that there was a minimum of 14 cm between cues. Each cue was presented when the mouse arrived 10 cm from its location, and disappeared once it was 4 cm behind the mouse. The portion of the maze where cues were presented (cue region) was 220 cm long, and after it the stem of the T-maze continued for another 80 cm where no cues were presented (delay region). At the end of the T-maze the mouse had to enter one of the arms, and full entry constituted a choice. Turning into the correct (more cues) side would elicit a water reward (6.4 uL), while an incorrect choice elicited a tone (pulsing 6 to 12 KHz tone for 1 s). At the time of reward or tone delivery, the visual environment froze for 1 s, and then disappeared for 2 s (after a successful trial) or 5 s (after a failed trial) before another trial was started.

### Pavlovian conditioning

After water restriction and handling, mice were habituated to head fixation for 2-3 sessions. Training consisted of 5 sessions (1 session/day); each session consisted of 50 reward deliveries (8 ul of water/reward). During training, each reward was preceded by a 2s tone that ended at the time of reward delivery. The time between a reward and the next tone delivery was sampled from an exponential distribution with a mean of 40s. The tone consisted of a sum of multiple sine waves with frequencies of 2, 4, 6, 8 and 16 Khz, and an amplitude of 70dB. All of the mice exhibited anticipatory licking by the end of the 5 days (increase in lick rate after tone presentation but before reward delivery). Some of the mice were previously trained for several days in a similar protocol where the tone amplitude was 60dB and the time between reward and subsequent tone was sampled from a uniform distribution between 5 and 15s; these mice did not exhibit anticipatory licking until trained in the final protocol. After training, RPE was assessed in a single test session that consisted of 64 trials; 50 of those trials were identical to the training trials (tone followed by reward), 7 trials were unexpected reward trials (reward delivery with no preceding tone) and 7 trials were unexpected omissions (tone not followed by reward). In all cases the intertrial interval was sampled from an exponential distribution with a mean of 40 s. Trial identity was sampled randomly with the following exceptions: 1- the first 5 trials were standard trials (tone+reward). 2- the first 2 non-standard trials were unexpected reward trials.

### Session and trial selection

We selected sessions and trials such that each recorded neuron would only appear in one session, and during which mice were engaged in the task. Our dataset contained one main imaging field/mouse, with the exception of three mice, in which we obtained two separate imaging fields at different depths. Thus, we analyzed 23 sessions from 20 mice (one session per imaging field). Sessions had at least 100 trials and mice performed at least 65% correct. Mice were between 3-6 months old during imaging and were trained for an average of 30 sessions before data collection (a range of 18-51 training sessions).

We removed a small fraction of trials in which mice were not engaged in the task, based on the following criteria: i) We calculated a smoothed performance measure by processing the binary trials success vector through a zero-phase filter composed of a 21 point centered Gaussian with std. dev.= 3. Trials where this measure was less than 0.5 were removed. ii) A sequence of 5 or more trials with the same choice and success rate equal or less than 20% was removed. iii) A sequence of 10 or more trials with the same choice was removed. The removed trials comprised 15% of trials per session on average. Most of these trials occurred close to the end of the session when the animals tended to exhibit decreased performance.

### Motion correction procedure

Deep brain imaging can be associated with spatially nonuniform fast motion (frame to frame), as well as spatially nonuniform slow drift of the field of view (over several minutes). To perform accurate motion correction despite the spatial non-uniformity, we divided the video into small regions (‘patches’) that had relatively uniform motion, and separately corrected the motion within each patch, as described below (schematic of procedure in Extended Data Fig. 3; example video before and after motion correction in Supplemental Video 1). Motion correction was performed on the red channel of the recording when available, otherwise it was performed on the green channel (n=6).

Before dividing the video into patches, we first performed rigid motion correction using a standard normalized cross-correlation method, to eliminate any spatially uniform motion (‘matchTemplate’ function in the openCV package in Python). This correction was performed on non-overlapping 50s video clips to eliminate concerns that slow drift over the course of minutes would degrade performance. The template for the cross-correlation was calculated by dividing each clip into non-overlapping sections of 100 frames, calculating the mean image of each section, and obtaining the median of the mean images. Before these motion correction steps, the video was pre-processed as follows: i- thresholded by subtracting a constant number and setting negative values to 0, such that the lower ~50% of pixels were 0, ii- used the openCV function ‘erode’ (with a scalar ‘1’ kernel), iii- convolved with a Gaussian (std. dev. = 2 pixels). Motion correction and template calculation were performed iteratively 10 times or until all absolute shifts were less than 1 pixel in both axes. Finally, the 50s clips had to be aligned to each other. This required generating a ‘master template’ for the entire video, and then using the same normalized cross-correlation procedure as before (‘matchTemplate’ function). The master template was calculated by taking the median of the templates of all clips.

The next step of motion correction involved compensating for spatially nonuniform, slow drift by estimating the drift in local patches. Patches were defined manually around neurons of interest to contain objects that drifted coherently (patch width ~80-160 pixels). In order to estimate the drift of each patch over time, we used a non-rigid image registration algorithm (demons algorithm, ‘imregdemons’ function in matlab). This algorithm outputs a pixel by pixel correction. However, directly applying this correction risks distorting the shape of the neurons or the amplitude of signals. Therefore, we applied a uniform correction for each patch, based on the average shift of all pixels in the patch (based on the demons output). We implemented the demons algorithm on the templates from the 50s clips described in the previous paragraph, again using the median of these templates as the ‘master template’. The registration and master template was computed iteratively 20 times, or until the increase in the average correlation between each corrected template and the overall template was less than the s.e.m. of these correlations. We found that the performance of the non-rigid registration improved if the templates were first processed through a local normalization procedure ^47^.

Finally, we performed standard rigid motion correction using the normalized cross-correlation method on each patch and each clip. We then repeated the rigid motion correction after taking a rolling mean of every two frames and downsampling the video by a factor of two. This increased signal strength; we used this downsampled video for subsequent analysis. After correcting for motion within clips, we had to correct across clips. To this end, we performed rigid motion correction on the clip templates. The motion correction code will be released on github upon acceptance of the manuscript.

### Calculation of ΔF/F from the motion-corrected images

The first step in calculating ΔF/F for each neuron was to define the neuron’s ROI, as well as the annulus around that ROI that would be used for neuropil correction ^48^^,^^49^. Each neuron’s ROI was defined manually using the mean and std projections of the movie as well as inspecting a movie that was downsampled by a factor of 5. An initial automatic annulus was generated by enlarging the borders of the ROI twice (by 5 um and 10 um); the annulus was the shape contained between the two enlarged borders, where we expect that observed activity would be due to neuropil but not the cell itself. Next, we manually reshaped the annulus region to avoid any visible dendrites, processes or cell bodies, while approximately maintaining its original area.

In order to correct for neuropil contamination, we subtracted a scaled version of the annulus fluorescence from the raw trace (*F_orr_(t)_=_ F_raw_(t) - ϒ·F_annulus_(t)*), where *F_raw_(t)* is the mean fluorescence in the neuron’s ROI at time *t*, *F_annulus_(t)* is the mean fluorescence in the corresponding annulus ROI at time *t*, and ϒ is the correction factor ^26^^,^^48^). A correction factor is needed since the neuropil activity is estimated in the imaged plane, while the actual contamination on the ROI is from out-of-plane fluorescence. The correction factor used was 0.58, which is in line with previously reported correction factors in GRIN lens imaging ^26^^,^^50^ and resulted in positive corrected traces. After neuropil subtraction, smoothing was performed by processing the corrected trace through a zero-phase filter using a 25 point centered Gaussian with 1.5 samples points std.

ΔF/F at time *t* was defined as (F(*t*)-F_0_(*t*))/F_0_(*t*), where F_0_(*t*) is the 8th percentile of the smoothed and neuropil corrected trace based on the preceding 60 seconds of recording.

### Selection of neurons in the dataset

Neurons were selected for analysis based on visual inspection of recording stability, using both the images as well as ΔF/F traces. Only neurons that were stable for at least 50 trials were included in the dataset. The full dataset comprised of n=303 neurons from n=20 mice. Of these, n=233 were considered to have a good fit by the encoding model described in the next section (>5% variance explained by the model during the cue period; reduced dataset). The full dataset was used in Fig. 2a, Fig. 4b, Extended Data Fig. 5, and Extended Data Fig. 7. For analyses where the specific output values of the encoding model were important, we used the reduced dataset composed of neurons for which the encoding model had a good fit (Fig 2c,d, Fig. 3, Fig. 4c, Extended Data Fig. 8, and Extended Data Fig. 10, Extended Data Fig. 11, and Extended Data Fig. 12).

### Encoding model

In order to quantify the contribution of behavioral variables to neural activity, we employed an encoding model, which was a multiple linear regression with the ΔF/F trace of each neuron as the dependent variable, and predictors derived from the behavioral variables as the independent variables (Fig. 2b). To derive the predictors, we divided the behavioral variables into 3 classes: “event” variables, “whole trial” variables, and “continuous” variables. “Event” variables (left and right cues, reward) were variables that occurred in discrete points in time. To derive the predictors for these variables, each event was convolved with a 7 degrees-of-freedom regression spline basis set with a 2 s duration, generated using the ‘bs’ package in R. “Whole-trial” variables (accuracy, previous reward) were variables whose value remained constant for an entire trial. These were coded as binary predictors, with a value of ‘1’ in all time points of trials where the animals received a reward (accuracy) or trials after receiving a reward (previous reward) and ‘0’ elsewhere. “Continuous” variables (position and kinematic variables) could change their value at every time point. In the case of kinematics, we included 3 “sub-variables” that were closely related to each other: velocity, acceleration, and view angle. Up to 3 predictors were generated per continuous variable (or sub-variable), by raising each variable to the 1^st^, 2^nd^ and 3^rd^ powers. The optimal number of predictors to use per continuous variable (for each neuron) was assessed by 5-fold cross-validation. (The reason that we used position along the maze as a continuous variable, rather than time in trial, was a previous study ^5^ which found that on a T-maze in which rats occasionally paused, DA activity seemed to be more closely related to position than time.)

The encoding model thus was:

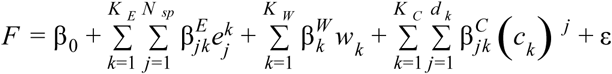

Where *F* is ΔF/F of a neuron, 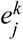 is is the j^th^ spline basis function convolved with the k^th^ event variable, *w_k_* is the predictor for the k^th^ whole-trial variable, *c_k_* is the k^th^ continuous variable, *K_E_*, *K_W_*, *K_C_* are the numbers of Event, Whole-trial, and Continuous variables correspondingly. *N_sp_* is the number of degrees of freedom for the spline (7 in all cases), *d_k_* is the maximal polynomial degree used for each continuous variable predictors, the β values are the regression coefficients for the different predictors, and ε is a Gaussian noise term. The β values were calculated using the least squares criterion after z-scoring the predictors (‘glmfit’ matlab function). This code will be released on github upon acceptance of the manuscript. Example single-trial fits for several cells are shown in Extended Data Fig. 4.

### Model comparison

We tested several behavioral variables on order to optimize the encoding model. The behavioral variables used in the final model (position, cues, kinematics, accuracy, previous reward) were those whose removal resulted in a significant degradation of the fit of the model prediction to the data across the population (Extended Data Fig. 5). Improved fits were assessed by comparing the R^2^ for each model (obtained with 5-fold crossvalidation) with a paired t-test across the population of neurons. We also considered other behavioral variables that did not improve the fit and therefore were not included in the final model (see Extended Data Fig. 5). The other variables that we considered are: *early and late cues*: a separate set of predictors was calculated for cues appearing in the 1^st^ half of the cue region and cues appearing in the 2^nd^ half. *#L - #R*: a predictor that at each timepoint takes the value of the current difference between left-and right-side cues that had appeared in the trial. |*#L - #R|* : a predictor that at each timepoint takes the absolute value of the current difference between left- and right-side cues that had appeared in the trial. *#L, #R*: two predictors that at each timepoint take the value of the current number of either left- or right-side cues that had appeared in the trial. *P(Reward on right) (nominal)*: a predictor that takes the current probability of the right side being rewarded based on the number of left- and right-side cues that had appeared in the trial and the sampling statistics of the cues. Given the Poisson distributions from which the cues were sampled (and ignoring the constraint of minimum distance between cues) this probability is given by the following logistic function: 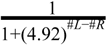, where *#L*, *#R* are the current counts of left- and right-sided cues respectively. The value of 4.92 is the ratio of Poisson means for high- and low-cue probability sides. *P(Reward on right) (empirical)*: a predictor that takes the current probability of the right side being rewarded based on the number of left- and right-side cues that had appeared in the trial, but instead of using the actual statistics of the cues, this probability was calculated using the psychometric curve of each mouse as the function that related the cue appearances to the probability of each side to be rewarded. Thus, this probability is given by : 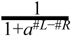 where the parameter *a* is estimated by fitting a logistic function to the psychometric curve of each mouse. *Difficulty of previous trial*: a predictor that is the final value of |*#L - #R|* from the previous trial. *Confirmatory/disconfirmatory cues*: Instead of dividing cues in left- and right-sided, cues are divided depending on whether they are confirming or disconfirming the current best estimate of the rewarded side. e.g. if the current count is 3 left-side cues and 1 right-side cue, if the next cue is a left-side cue it is confirmatory, and if it is a right-side cue it is disconfirmatory (in case of an even count the next cue is considered confirmatory).

### Calculation of the relative contributions of behavioral variables to neural activity

We quantified the relative contribution of each behavioral variable to neural activity (Fig. 2c,d) by determining how the performance of the encoding model declined when each variable was excluded from the model. We predicted neural activity with all variables (“full model”) or by excluding one of the variables (“partial model”), in either case with 5-fold cross-validation. The relative contribution of each behavioral variable was calculated by comparing the variance explained of the partial model to the variance explained of the full model. In the case of the cue period, in which five behavioral variables, relative contribution of each variable was defined as 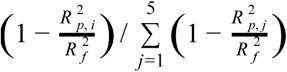 where *R*^2^_*p, i*_ is the variance explained of the partial model that excludes the *i^th^* variable and *R*^2^_*f*_ is that of the full model. In the case of the outcome period, two event variables were considered: time of reward and time of outcome (reward or tone delivery). The relative contribution of reward was calculated by comparing the variance explained of a partial model with only the time of outcome, compared to a full model that had both time of reward and time of outcome as event predictors, 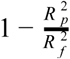. This allowed us to identified variance in the neural activity that could be attributed to reward rather than simply reaching the end of the maze. When fitting the model for the calculation of the relative contribution for reward, we matched the overall effects of correct and incorrect trials by overweighting incorrect trials for each crossvalidation fold. Negative relative contributions were clamped at 0 (this occurs when the *R^2^* of the full model is lower than that of the partial model, due to introduction of noise by the excluded variable).

We used two approaches to exclude variables from the full model and calculate variance explained by the partial model. In the first approach, the partial model was equivalent to the full model, except that the β values of the predictors of the excluded variable were set to zero (“no refitting”). In the second approach, we calculated new β values by re-running the regression without the predictors of the excluded variable (“refitting”). Both approaches to exclude variables produced comparable results; the “no refitting” approach was used to generate the main figures, while comparison with the “refitting” approach is shown in Extended Data Fig. 12.

To determine if the contribution of a behavioral variable was statistically significant for each neuron (Fig. 2a; Extended Data Fig. 7), we first calculated the F-statistic of the nested model comparison test where the reduced model was the model without that behavioral variable included. We then proceeded to calculate the same statistic on 1000 instances of shuffled data, where shuffling was performed on non-overlapping 3s bins (to maintain the autocorrelation of the signal). The p-value used for significance was obtained by comparing the value of the original F-statistic to the shuffle distribution, using the Holm-Bonferroni correction to account for the number of behavioral variables tested for each neuron; the threshold for significance was a p-value of 0.01 after correction.

### Clustering analysis

To identify functional clusters of neurons (Fig. 3a), we used a clustering procedure based on a Gaussian mixture model (GMM) that was applied on the matrix of contributions of behavioral variables to the neural activity. To do that, we used the ‘fitgmdist’ function in Matlab (Mathworks, Inc) with 1000 maximum iterations, 0.35 regularization value, 100 replicates, and the covariance matrix constrained to diagonal. This produces a Gaussian mixture model where the major axes of the Gaussians are parallel to the axes of the feature space, which enables flexibility beyond that of the k-means algorithm while still maintaining a relatively small number of parameters to be fitted.

To test the fit of the clustering model (Fig. 3b), we shuffled 10,000 times the relative contribution values both across behavioral variables (Fig. 3b, top) and across neurons (Fig. 3b, bottom; the contributions for the cue period variables were re-normalized per neuron after shuffling). After each shuffling iteration, we repeated the clustering and recalculated the log-likelihood of the clustering model. The distribution of log-likelihood values for shuffled data was then compared to the log-likelihood of the clustering model on the real data.

The BIC score was used to select the number of clusters. It is a penalized likelihood term defined as *2(NlogL)* + *Mlog(n)*, where *NlogL* is the negative log-likelihood of the data, *M* is the number of parameters of the GMM, and *n* is the number of observations. The first term rewards model with good fit, while the second term penalizes more complex models. The BIC score was calculated by the ‘fitgmdist’ function.

### Quantification of reward prediction error signals with d'

In figure 5, the strength of modulation of reward responses by reward expectation was calculated using the d’ measure as follows: 1- We divided rewarded trials into trials with either high reward expectation (HRE) or low reward expectation (LRE). For the pavlovian conditioning experiments, HRE trials were those where reward delivery was preceded by a tone, and LRE trials were those where reward delivery was not preceded by a tone. For the virtual reality experiments, trials were divided in two different ways: for the trial difficulty criterion, we ranked trials according to the strength of the evidence (absolute value of the difference between the total number of right- and left-sided cues). The top half of those trials (strong evidence) were considered HRE trials and the bottom half (weak evidence) were considered LRE trials. For the previous outcome criterion, previously rewarded trials were HRE trials and previously unrewarded trials were LRE trials. 2- We calculated the average reward response in each trial by averaging activity in the first 2 s following reward delivery and subtracting from that the average activity in the 1 s preceding reward delivery. 3- The d’ for the reward responses for HRE and LRE trials was calculated as follows:

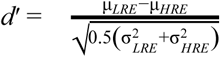

where μ and σ^2^ are the mean and variance of the distribution of reward responses for the denoted trial group. Thus, positive d’ values indicate activity consistent with a reward prediction error signal (stronger reward response for low reward expectation trials).

### Histology

After completion of behavioral experiments, mice were perfused with 4% PFA in PBS, and then brains were removed and postfixed in 4% PFA for 24 additional hours before transferring to 30% sucrose in PBS. After post-fixing, 40 micron sections were made with either a microtome (American Optical 860) or cryostat (Leica CM3050 S). Brain sections were washed with PBST (Phosphate buffered saline with 0.4% Triton x-100) for 30 min, and then placed in blocking buffer (10 ml PBST + 0.2 ml normal donkey serum + 0.1 g bovine serum albumin (sigma A7906-100G) for 1 hour. Sections were incubated overnight at 4° C in primary antibodies for TH (TH Ab; Aves labs, E.C. 1.14.16.2, chicken polyclonal anti-peptide antibody mixture, 1:1000 dilution) and GFP (Molecular probes G10362, rabbit monoclonal, 1:1000 dilution). Sections were then washed with PBST for 30 min, then incubated for 1 hour at room temperature in Alexa fluor 647 (Jackson ImmunoResearch Donkey-anti-chicken, 1:1000 dilution) and Donkey anti-rabbit Alexa fluor 488 (Jackson ImmunoResearch, 711-545-152, 1:1000 dilution). Following PBST washes, sections were mounted in 1:2500 DAPI in Fluoromount-G. Whole sections were imaged with a Nikon Ti2000E microscope.

### Estimation of the neurons’ location

In order to investigate the relationship between the activity of the neurons and their location in the VTA (Fig. 3d), we estimated each neuron’s location by combining information about the position of the GRIN lens from histology with the location of the imaged neurons within the field of view. Histological slices stained for Tyrosine hydroxylase (TH) featuring the tract left by the GRIN lens (Extended Data Fig. 9) were manually scaled and rotated to fit images taken from a mouse atlas ^51^ using the VTA, SNc and cerebral peduncle as primary markers. The center of the bottom of the lesion was used as a proxy for the center of the lens, and its location was measured using the atlas coordinates. We estimate the error of our determination of the location of the lens as follows: for the M/L axis, for several mice, we independently repeated the procedure using different slices. The difference in the obtained estimate was +/-50 um. Thus, we consider that to be our error. For the A/P axis, the resolution depended on comparing coronal slices to the atlas images, which are provided with a sampling of 115 um on average. Thus, we estimate our error to be within two such images, or +/-115 um.

To calibrate the diameter of each field of view, we first estimated the average size of DA neurons in the VTA. To do that, an Ai148xDAT::cre mouse was sacrificed. The brain was extracted and then immersed in ice-cold NMDG ACSF for 2 min. Afterward, coronal slices (300 um) were sectioned using a vibratome (VT1200S, Leica) and then incubated in NMDG ACSF at 34° for 15 min. Then the slices were incubated in PFA for 5 min, transferred to a slide, and immediately imaged under the 2-photon microscope. We manually traced the ROIs of the imaged neurons, and then fit an ellipse to each ROI using the direct method ^52^. The average values for the minor and major axes of the ellipse was 9 and 15 um respectively. For each field of view imaged during the experiment, we repeated this procedure and calculated the average minor and major axes of the neurons’ ROIs. The size of each field of view was calibrated such that we minimized the mean square difference between the measured ellipse values and the average ones previously derived. The center of mass of each ROI was used as the marker for the neuron location within the field of view. The absolute location of the neuron was the vector sum of its distance from the lens center in the field of view to the measured location of the lens center in atlas coordinates. These estimates were used in Figs. 1, 3 & 4 and Extended Data Figs. 10 & 11.

The relative concentration across the A/P or M/L axis of neurons belonging to a given cluster (Fig. 3e) was calculated as follows. First, the concentration of neurons belonging to a cluster was estimated using Gaussian kernel smoothing via the ‘ksdensity’ function in Matlab with a bandwidth of 50 um applied only on these neurons. Second, the relative concentration for each cluster was calculated as the concentration per cluster divided by the sum of concentrations calculated for all clusters. To calculate the 95% confidence intervals of the relative concentrations (Fig. 3e, dashed lines), we ran 10000 iterations where in each we randomized the cluster identities of the neurons and then proceeded to calculate the relative concentrations of each cluster as above. For each point in the A/P or M/L axis, the edges of confidence interval were the 2.5 and 97.5 percentiles of the distribution of concentrations calculated from the shuffled data.

### Signal and noise correlations

To investigate how the correlations between pairs of neurons were spatially organized in the VTA, we calculated signal and noise correlations for all pairs of neurons that were simultaneously recorded (Fig. 4c). The signal correlation between a pair of neurons was calculated by correlating the predictions of the encoding model for both neurons in the cue period or outcome period. The noise correlation was the correlation between the residuals for each neuron pair. We also used an alternative method for estimating the noise correlations ^30^^,^^31^ (Extended Data Fig. 11). The alternative noise correlation estimate between a pair of neurons (*i*,*j*) was calculated as follows: we first fit an augmented encoding model for neuron *i* which had as an additional predictor the activity of neuron *j*; we then calculated the normalized improvement in the fit using 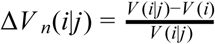 where *V* (*i*|*j*), *V* (*i*) are the variances explained by the augmented and original (behavioral-only) encoding models respectively for neuron *i*. We repeated this procedure for neuron *j* and obtained Δ*V_n_* (*j*|*i*) . The noise correlation estimate was the mean of the two Δ*V_n_* values.

### *Ex vivo* recordings to compare GCaMP6f fluorescence with activity in DA neurons

In order to compare GCaMP6f fluorescence with spike times in DA neurons (Extended Data Fig 2), we performed *ex vivo* slice imaging and electrophysiolgical recordings in Ai 148xDAT:: Cre mice. Mice were anesthetized with an i.p. injection of Euthasol (0.06ml/30g) and decapitated. After extraction, the brain was immersed in ice-cold carbogenated NMDG ACSF (92 mM NMDG, 2.5 mM KCl, 1.25 mM NaH2PO4, 30 mM NaHCO3, 20 mM HEPES, 25 mM glucose, 2 mM thiourea, 5 mM Na-ascorbate, 3 mM Na-pyruvate, 0.5 mM CaCl2·4H2O, 10 mM MgSO4·7H2O, and 12 mM N-Acetyl-L-cysteine) for 2 minutes. The pH was adjusted to 7.3-7.4. Afterwards coronal slices (300um) were sectioned using a vibratome (VT1200s, Leica) and then incubated in NMDG ACSF at 34°C for 15 minutes. Slices were then transferred into a holding solution of HEPES ACSF (92 mM NaCl, 2.5 mM KCl, 1.25 mM NaH2PO4, 30 mM NaHCO3, 20 mM HEPES, 25 mM glucose, 2 mM thiourea, 5 mM Na-ascorbate, 3 mM Na-pyruvate, 2 mM CaCl2·4H2O, 2 mM MgSO4·7H2O and 12 mM N-Acetyl-l-cysteine, bubbled at room temperature with 95% 02/ 5% CO2) for at least 45 mins until recordings were performed.

During cell-attached recordings, slices were perfused with a recording ACSF solution (120 mM NaCl, 3.5 mM KCl, 1.25 mM NaH2PO4, 26 mM NaHCO3, 1.3 mM MgCl2, 2 mM CaCl2 and 11 mM D-(+)-glucose, continuously bubbled with 95% O2/5% CO2). Cell-attached recordings were performed using a Multiclamp 700B (Molecular Devices, Sunnyvale, CA) using pipettes with a resistance of 4-6 MOhm filled with an internal solution identical to the recording ACSF. Infrared differential interference contrast–enhanced visual guidance was used to select neurons that were 3–4 cell layers below the surface of the slices, which were held at room temperature while the recording solution was delivered to slices via superfusion driven by peristaltic pump. Cell-attached recordings were collected once a seal (200 MOhm to >5 GOhm) between the recording pipette and the cell membrane was obtained. To generate bursts in cells that did not exhibit spontaneous bursting activity, a second glass pipette filled with recording ACSF containing 20 µM NMDA was placed above the recorded cell. Slight positive pressure (~12 psi) was briefly applied (100-250 ms) to generate bursting activity in the recorded cell. During bursts, spikes typically exhibited a gradual reduction in amplitude as observed previously ^53^. Action potential currents were recorded in voltage-clamp mode with voltage clamped at 0 mV, which maintained an average holding current of 0 pA. Cell-attached currents were low-pass filtered at 1 kHz and digitized and stored at 10 kHz (Clampex 9; MDS Analytical Technologies). All experiments were completed within 4 hours after slicing the brain. Fluorescence was imaged using a CMOS camera (ORCA-Flash 2.8, Hamamatsu) at 30Hz using a GFP filter cube set (exciter ET470/40x, dichroic T495LP, emitter ET525/50m).

